# Evaluation of the accuracy of bacterial genome reconstruction with Oxford Nanopore R10.4.1 long-read-only sequencing

**DOI:** 10.1101/2024.01.12.575342

**Authors:** Nicholas D Sanderson, Katie Hopkins, Matthew Colpus, Melody Parker, Sam Lipworth, Derrick Crook, Nicole Stoesser

## Abstract

2.

Whole genome reconstruction of bacterial pathogens has become an import tool for tracking antimicrobial resistance spread, however accurate and complete assemblies have only been achievable using hybrid long and short-read sequencing. We have previously found the Oxford Nanopore Technologies (ONT) R10.4/kit12 flowcells produced improved assemblies over the R9.4.1/kit10, however they contained too many errors compared to hybrid Illumina-ONT assemblies. ONT have since released the R10.4.1/kit12 flowcells that promises greater accuracy and yield. They have also released newly trained basecallers using native bacterial DNA containing methylation sites intended to fix systematic errors, specifically Adenosine (A) to Guanine (G) and Cytosine (C) to Thymine (T) substitutions. ONT have recommended the use of Bovine Serum Albumin (BSA) during library preparation to improve sequencing yield and accuracy. To evaluate these improvements, we sequenced DNA extracts from four commonly studied bacterial pathogens, namely *Escherichia coli*, *Klebsiella pneumoniae*, *Pseudomonas aeruginosa* and *Staphylococcus aureus*, as well as 12 disparate *E. coli* clinical samples from different phylogroups and sequence types. These were all sequenced with and without BSA. These sequences were *de novo* assembled and compared against Illumina-corrected reference genomes. Here we have found the nanopore long read-only R10.4.1 (kit14) assemblies with basecallers trained using native bacterial methylated DNA produce accurate assemblies from 40x depth or higher, sufficient to be cost-effective compared to hybrid long-read (ONT) and short-read (Illumina) sequencing.

**Impact statement:** Currently, the best method of building accurate and complete bacterial genome assemblies is to create a hybrid assembly; combining both long and short DNA sequencing reads. Short reads are more accurate, but can be difficult to assemble into a complete genome. Long reads are generally less accurate, but easier to reconstruct into a complete genome. By combining long and short reads, we get both accuracy and reconstructive power. However, this also involves higher costs and more labour than using a single sequencing platform. In this study, we compare long read only assemblies from Oxford Nanopore Technology’s newest iteration of improvements in both chemistry and software to hybrid Illumina-Nanopore assemblies. We sequenced four bacterial pathogens with published reference genomes (*Staphylococcus aureus, Klebsiella Pneumoniae, Pseudomonas Aeruginosa*, and *Escherichia Coli*) and twelve bloodstream associated *E. coli*, and show that assemblies from the newest technology are not only an improvement on the previous iteration, but are able to compete with hybrid Illumina-Nanopore assemblies in their quality, providing a step towards bacterial genome assembly using a single sequencing platform.

**Data summary:** **The authors confirm all supporting data, code and protocols have been provided within the article, through supplementary data files, or in publicly accessible repositories.**

Nanopore and Illumina fastq data are available in the ENA under project accession: PRJEB51164.

Assemblies have been made available at: https://figshare.com/articles/dataset/R10_4_1_KIT14_comparison_assemblies/2497 2954

Code and analysis outputs are available at:

https://gitlab.com/ModernisingMedicalMicrobiology/assembly_comparison

## 5. Introduction

Generating high quality whole genome *de novo* assemblies for bacterial pathogens has become an important tool supporting diagnostic, infection prevention control and public health initiatives, improving our understanding of antimicrobial resistance and pathogen epidemiology[1]. The introduction of longer sequencing reads from PacBio and Oxford Nanopore Technologies (ONT) has enabled the reconstruction of previously difficult-to-assemble genomes. Previously we have found the most cost-effective method of assembling high-accuracy bacterial genomes is to combine long R9.4.1/kit10 ONT reads with high-accuracy short Illumina reads[2]. Generating these high-quality assemblies without the need for Illumina sequencing could reduce costs and laboratory time, and this would enable more cost-effective, faster, and higher-throughput sequencing.

ONT have iteratively updated their flowcell technology and chemistry from R9.4.1/kit10 through R10.3/kit12 to R10.4/kit12 and most recently R10.4.1/kit14; older flowcells and chemistries have been phased out. Further changes to ONT workflows have included an increase in sampling rate on the nanopore devices from 4k samples per second to 5kHz, which generates more data points for basecalling and thereby improves accuracy, according to ONT [3]. Users of R10.4 have reported cytosine to thymine (C-to-T) and guanine to adenine (G-to-A) errors, hypothesised to be caused by methylated sites confusing the basecalling models. ONT have released a research basecalling model (rerio) trained with native bacterial DNA and designed to fix errors associated with methylated sites which also takes advantage of the increased sampling rate. It is now also recommended to use bovine serum albumin (BSA) which is added during library preparation to improve sequencing yield and quality by blocking non-specific binding sites.

We have previously compared ONT-only assemblies using R10.3 or R10.4 sequencing data to the R9.4.1//Illumina hybrid assemblies[4]. We looked at four common, diverse human pathogenic reference strains representing a range of %GC content, namely, *Staphylococcus aureus* MRSA252*, Klebsiella pneumoniae* MGH78578*, Pseudomonas aeruginosa* PAO1, and *Escherichia coli* CFT073. R10.4-only assemblies (using superior accuracy [sup] basecalling) looked promising in terms of genome reconstruction, error rates and gene recovery, but generally required deeper long-read sequencing depths than for a hybrid assembly, rendering the latter more cost-effective and high-throughput if access to both ONT and Illumina sequencing modalities was available[4]. Smaller plasmids (<5kb) were also less effectively recovered as part of long-read only sequencing. These findings have been consistent with other more recent studies[5, 6].

As an update to our previous evaluation, here we compare the accuracy of assemblies from nanopore-only R10.4.1 sequencing with our previous hybrid R9.4.1/Illumina assemblies for the same four species references. Given that *E. coli* was the most challenging species to assemble previously, we also evaluate the accuracy of R10.4.1-only assemblies for 12 *E. coli* strains representing major phylogroups of the species compared with R9.4.1/Illumina hybrids created for these *E. coli* strains. As part of this second evaluation, we characterise the impact of BSA on sequencing output, the impact of increased sampling rate and the newer basecalling models on read and assembly accuracy, and estimate current sequencing costs per isolate using R10.4.1 long-read only assembly.

## 6. Methods

### 6.1 Bacterial isolates and DNA extraction

DNA was extracted from laboratory stocks of four bacterial reference strains: *P. aeruginosa* PAO1*, S. aureus* MRSA252*, K. pneumoniae* MGH78578*, E. coli* CFT073, and twelve bloodstream-associated *E. coli* isolates, for which we had existing Illumina data[7] and which were chosen to reflect the genetic diversity within the species based on *in silico* phylotyping[8]. Stocks were stored at -80°C in nutrient broth with 10% glycerol. Stocks were cultured on Columbia Blood Agar overnight at 37°C. One colony was selected and sub-cultured for DNA extraction using the QIAGEN Genomic Tip 100/G kit. Extracts were stored in elution buffer at 4°C for the duration of the study. DNA quality was assessed with the Qubit Fluorometer and TapeStation immediately after extraction, and periodically prior to sequencing thereafter.

### 6.2 Nanopore sequencing

All sequencing occurred on an ONT GridION device (Minknow version 23.04.5) using a 5kHz sampling rate with R10.4.1 flow cells and sup base calling enabled. Sequencing libraries were created using the Rapid Barcoding (RBK114.96) kit, and each run was set for 72 hours.

For the four reference strains, we followed the recommendation to use BSA (50mg/mL, Thermo Fisher Scientific, AM2616) as part of library preparation and created a multiplexed library using the four reference strains which was run on a single R10.4.1 flowcell. The yield for *K. pneumoniae* on this run was too low (10.8 Mb only), so this extract was sequenced again as a single sample on a second R10.4.1 flowcell. To specifically assess the impact of adding BSA to the flowcell priming mix, a multiplexed library of the twelve *E. coli* isolates was created, and split between two R10.4.1 flowcells. One of these flowcells included the addition of BSA to the flowcell priming mix, and the other did not Sampling rates, basecalling and runtimes were as above.

R9.4.1 data were generated previously[4].

### 6.3 Illumina sequencing

Illumina sequencing data for the reference strains were generated previously as part of [4], and for the 12 *E. coli* isolates as part of [7]

A summary of the experimental workflow is shown in Supplemental Figure S1.

### 6.4 Hybrid Assemblies

#### 6.4.1 Reference strains

The four reference strains were hybrid assembled with nanopore and Illumina sequences using Unicycler (v0.5.0, default settings) [9]. Hybrid assemblies were generated with both R9.4.1 and R10.4.1 nanopore data were for comparison. The nanopore reads were subsampled to specific depths but the complete quantity of Illumina reads was always used.

### 6.4.2 *E. coli* isolates

For the 12 clinical *E. coli* isolates where no gold standard reference genomes were available, hybrid Illumina/nanopore assemblies were generated (using R9.4.1 data) with Unicycler (v0.5.0, default settings, except G_117, E_8318 where flye assemblies were used instead of miniasm within unicycler) to act as references[9]. These were compared to R10.4.1 long-read only assemblies, generated using the optimal strategy developed on the four mixed species reference strains, namely Dorado/Rerio basecalling + Flye assembly with a single round of Medaka polishing.

### 6.5 Data analysis

An analysis workflow was written in nextflow and is publicly available in gitlab (https://gitlab.com/ModernisingMedicalMicrobiology/assembly_comparison). Per read accuracy, subsampling, assembly, contig reference comparison and gene recovery metrics are all described previously[4]. Briefly, nanopore reads were subsampled (20 to 100x depths at 10x intervals) using Rasusa (v0.6.1)[10], then assembled with Flye[11] (using the --meta and –nano-hq parameters, version 2.9-b1768) along with the un-subsampled reads (represented in figures as “0”). Medaka (1.6.0; default settings; https://github.com/nanoporetech/medaka) was used to polish the assembled contigs from Flye. Only one polishing iteration was used as further polishing cycles did not improve error rates and used significant compute resource (Supplemental Figure S2 [also shown in our previous evaluation[4]]).

Given that deposited reference sequences may contain uncertainties (e.g. R, Y, W or K IUPAC nucleotide codes) and long-term storage of reference sequences may lead to minor genuine genetic changes, reference genome sequences from *E. coli* CFT073 (AE014075.1), *K. pneumoniae* MGH78578 (CP000647.1), *P. aeruginosa* PAO1 (NC_002516.2), *S. aureus* MRSA252 (NC_002952.2) were corrected with Illumina sequence data from [8] using the SNIPPY workflow (version 4.6.0) (https://github.com/tseemann/snippy). For empirical read accuracy, reads were mapped to these Illumina-corrected reference sequences with minimap2 [12] (version 2.22-r1101) and samtools[13], and the percentage identity was calculated from the query distance (NM tag) in the bam file over the query length, multiplied by 100. Empirical Q-scores were not shown due to the number of reads with no errors resulting in infinite values that skewed the results.

### 6.5.1 Basecalling

Raw R9.4.1 nanopore data from [8] was re-basecalled using guppy version 5.0.12+eb1 a981 with the dna_r9.4.1_e8.1_sup.cfg model. R10.4.1 nanopore raw data was basecalled using Dorado version 0.4.0+0aaf16d and either dna_r10.4.1_e8.2_400bps_5khz_sup.cfg (5k RAPID) or res_dna_r10.4.1_e8.2_400bps_sup@2023-09-22_bacterial-methylation (rerio) models.

### 6.5.2 Assembly error profiling

The error types from the assembled contigs were characterized using the output from DNADIFF (Mummer3 package [14]) and summarised based on type insertion or deletion (INDEL) and single nucleotide differences (SNV).

### 6.6 Statistical analysis and visualisations

Mann-Whitney-Wilcoxon (two-sample Wilcoxon) tests were used to evaluate the statistical significance of pairwise differences in continuous non-normal variables such as read length and accuracy for each sequencing modality. Statistical analysis and data visualisations were carried out in R version (version 4.3.0) using the ggpubr and ggplot packages, and Supplementary figure S 1 was generated in Biorender (www.biorender.com).

## 7. Results

### 7.1 Four bacterial reference strains - Sequencing yield and read length distributions

The R9.4.1 flowcell generated more data than the combined R10.4.1 flowcells after running for 72 hours, (10.5Gb versus 5.2Gb, Supplementary figure S 3 and Table 1). The data generated for the two R10.4.1 results and plotted originate from the same combined read yields from two flowcells (as *K. pneumoniae* was re-sequenced separately) but differ by basecalling method (Dorado/SUP versus Dorado/rerio methods), resulting in slightly different total base outputs (Supplementary figure S 3).

**Table 1.** **Total bases, and median (IQR) read lengths excluding the unclassified barcode, per flowcell and per species isolate on each flowcell.**

| Run type | Flow cell bases | Species | Q1 length | Median length | Q3 length | Isolate bases |
| --- | --- | --- | --- | --- | --- | --- |
| r9.4.1 4k BSA- Rapid Guppy SUP | 10,538,807,369 | *E. coli* | 1,565.75 | 4,058.00 | 8,345.25 | 2,242,222,750 |
|  |  | *K. pneumoniae* | 2,729.00 | 5,245.00 | 12,007.50 | 3,458,646,526 |
|  |  | *P. aeruginosa* | 2,956.75 | 6,869.00 | 15,058.00 | 4,138,688,286 |
|  |  | *S. aureus* | 4,195.50 | 10,920.50 | 24,525.00 | 699,249,807 |
| r10.4.1 5k BSA+ Rapid Dorado SUP | 5,238,710,880 | *E. coli* | 389.75 | 1,914.50 | 8,283.50 | 2,617,431,144 |
|  |  | *K. pneumoniae* | 410.50 | 2,048.00 | 6,810.50 | 417,791,092 |
|  |  | *P. aeruginosa* | 264.00 | 533.00 | 2,848.75 | 1,660,043,947 |
|  |  | *S. aureus* | 647.50 | 3,675.00 | 10,926.00 | 543,444,697 |
| r10.4.1 5k BSA+ Rapid Dorado rerio | 5,187,428,875 | *E. coli* | 379.25 | 1,906.00 | 8,060.25 | 2,584,825,627 |
|  |  | *K. pneumoniae* | 486.75 | 2,326.00 | 7,631.50 | 417,332,669 |
|  |  | *P. aeruginosa* | 266.00 | 507.00 | 2,972.75 | 1,654,741,681 |
|  |  | *S. aureus* | 535.00 | 3,294.00 | 10,844.50 | 530,528,898 |

Median read lengths varied between species for each run type with R9.4.1 having the highest median and IQR lengths across all species (median read length for R9.4.1: 6.5kb [IQR: 2.68-15.2kb] versus for R10.4.1: 1.7 kb [IQR: 0.34 -7.73 kb]; p<0.001, two-sample Wilcoxon) (Supplementary figure S 4).

### 7.2 Four bacterial reference strains - Empirical read accuracy

The empirical accuracy of the raw reads was determined from alignments to the Illumina-corrected reference genomes. Of the long-read methods, the R9.4.1 reads had the lowest accuracy (median: 96.8% [IQR: 95.9-97.4%]) compared with R10.4.1 reads (median: 98.8%, [IQR: 98.1-99.2%], p <0.001, two-sample Wilcoxon) 94%; Figure 1). The modal accuracies ranged from 97% to 97.5% for R9.4.1 and increased to >99% for the R10.4.1 reads. Reads basecalled using Dorado and the rerio research model showed marginally higher read-level accuracies compared to the Dorado/SUP basecalled data (median: 98.7% [IQR: 97.9-99.1%] vs median: 98.8%, [IQR: 98.1-99.2%], p<0.001, two-sample Wilcoxon).

**Figure 1.**
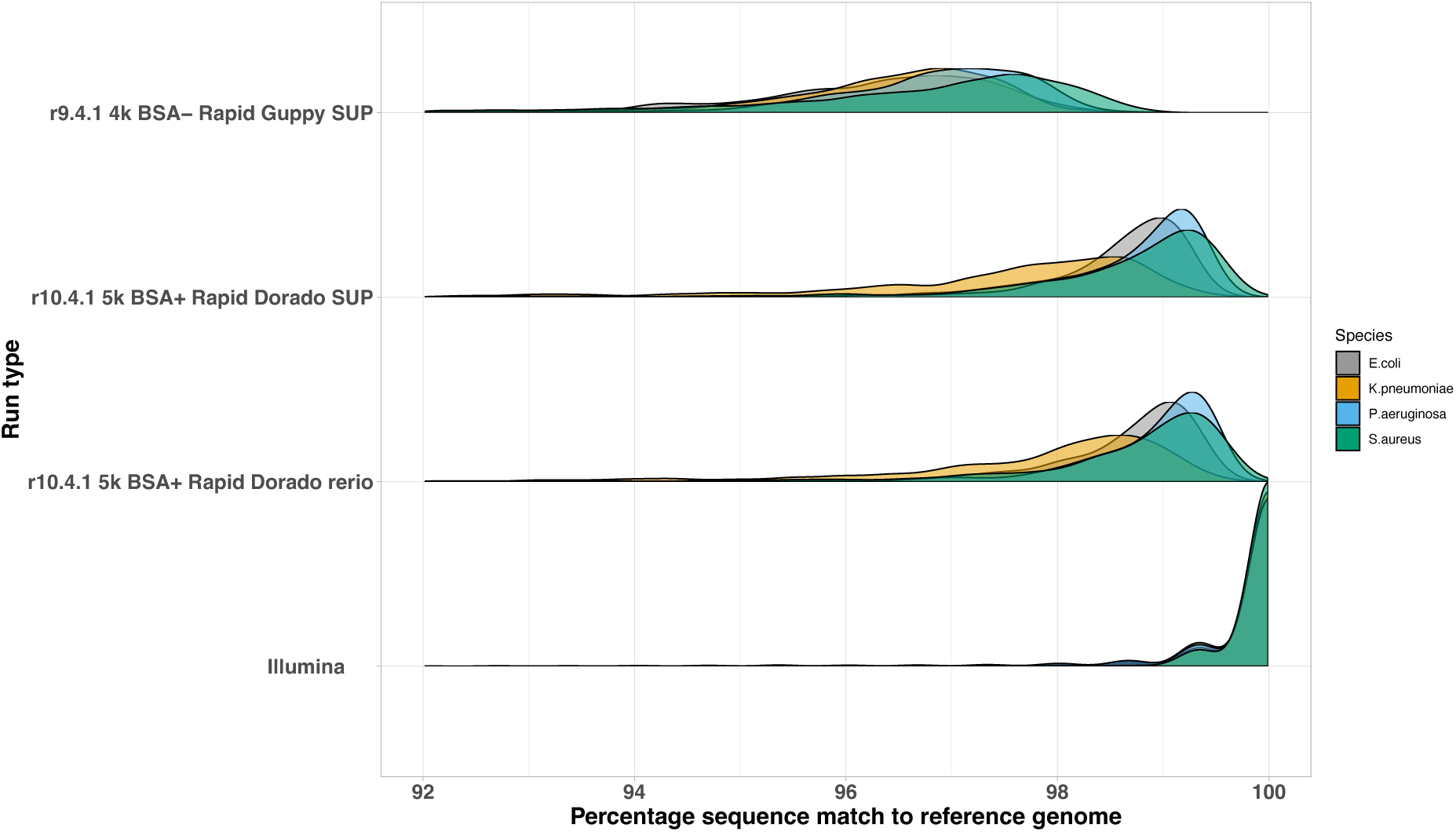
**Distributions of read percentage match to the reference genome for sequenced reference strains by sequencing run type and bacterial species.**

### 7.3 Four bacterial reference strains - Assembly chromosome and plasmid recovery

The nanopore-only R10.4.1 sequences were assembled with Flye and polished with medaka, the nanopore and Illumina hybrid assemblies were assembled with Unicycler; hybrid assemblies were generated with both R9.4.1 and R10.4.1 nanopore data for comparison. The nanopore-only (Flye and medaka) and R10.4.1+Illumina hybrid (Unicycler) assemblies all fully assembled the reference chromosome sequences into single contigs at all subsampled sequencing depths, Figure **2**. The R9.4.1+Illumina hybrid (Unicycler) approach reconstructed the *E. coli* chromosome as two contigs (as opposed to a fully resolved, single contig) at 20 and 30x subsampled depths but completely resolved the reference strain chromosomes at all other depths (Figure **2**).

**Figure 2.**
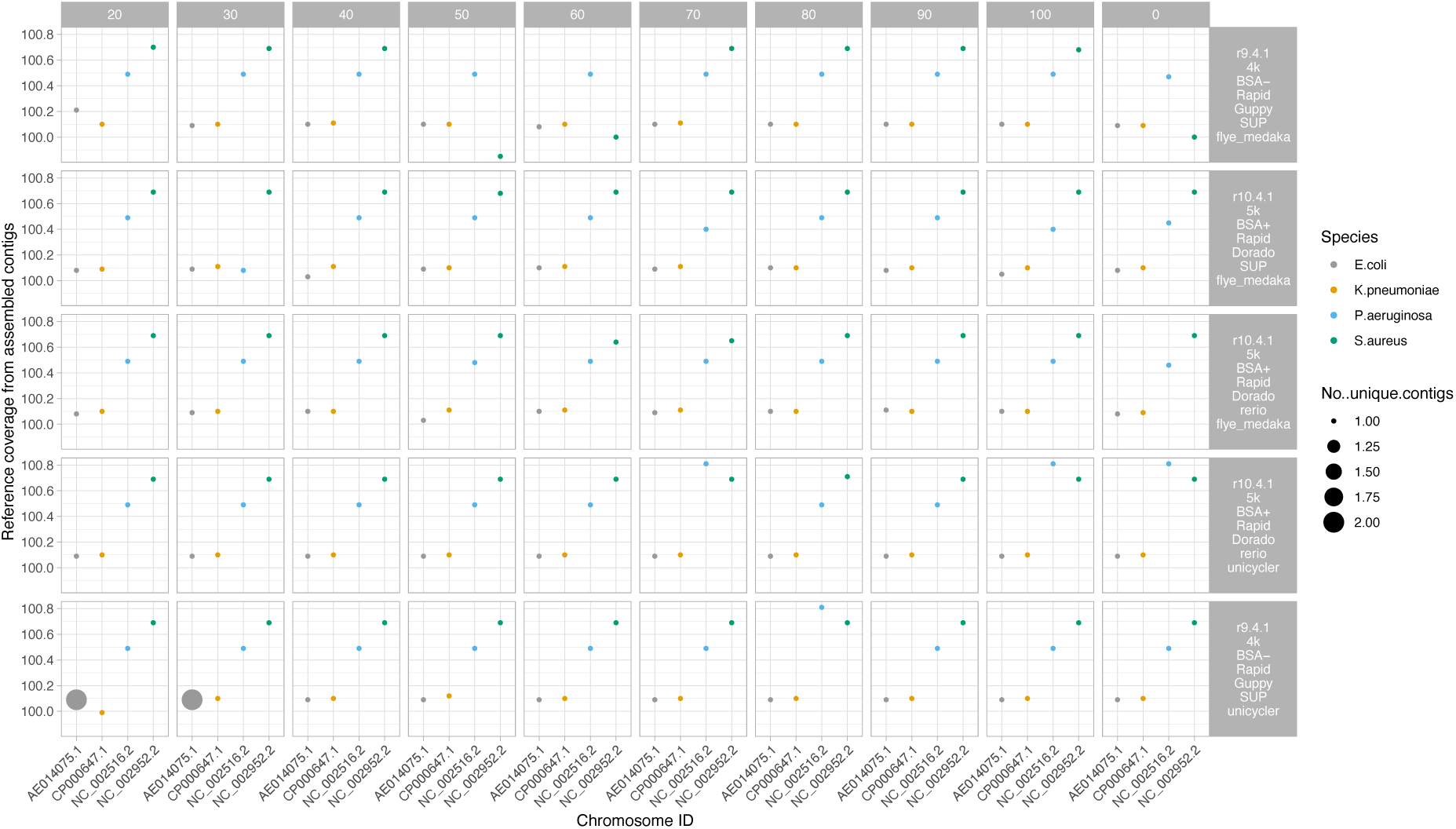
**Percentage of reference strain chromosomes recovered by run type and species. Size of dots represent the number of unique contigs recovered. Faceted columns represent subsampling to a specified depth (x coverage), with 0 the full number of reads from the run used. Faceted rows represent run type, including the flowcell cell used (R9.4.1 or R10.4.1), sampling rate (4k or 5k), if BSA was used (BSA+ or BSA-), the basecaller used (Guppy or dorado), the basecalling model used (SUP or rerio), and the assembly strategy used (Flye+Medaka or Unicyler) Unicycler assemblies represent nanopore-Illumina hybrid assemblies.**

Plasmid recovery for most long-read only assemblies was affected by either higher sequencing depth (i.e. impaired at >100x depth for R9.4.1 guppy/sup basecalled and R10.4.1 dorado/sup basecalled data) or at <40x sequencing depth (for R10.4.1 dorado/rerio basecalled data), but was otherwise excellent (Figure **3**). Both R9.4.1+Illumina and R10.4.1+Illumina hybrid assemblies also resulted in excellent plasmid recovery with long read sequencing depths of 20x-100x.

**Figure 3.**
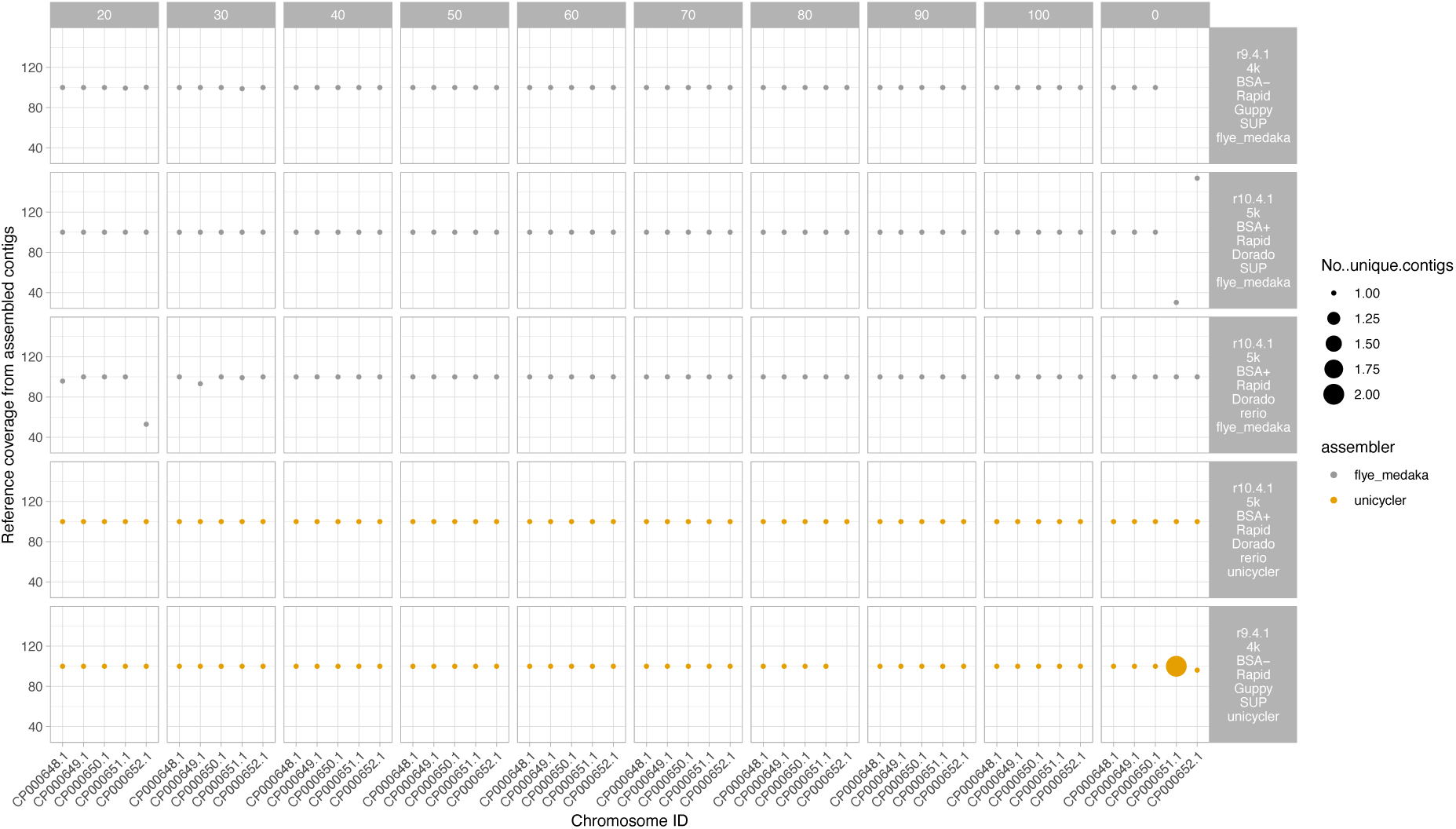
**Percentage of reference strain *K. pneumoniae* plasmid sequences recovered by run type. Size of dots represent the number of unique contigs. Faceted columns represent subsampling to a specified depth (x coverage), with 0 the full number of reads from the run used. Faceted rows represent run type, including the flowcell cell used (R9.4.1 or R10.4.1), sampling rate (4k or 5k), if BSA was used (BSA+ or BSA-), the basecaller used (Guppy or dorado), the basecalling model used (SUP or rerio), and the assembly strategy used (Flye+Medaka or Unicycler) Unicycler assemblies represent nanopore-Illumina hybrid assemblies.**

### 7.4 Four bacterial reference strains - Assembly accuracy

Comparing the assembled contigs to the Illumina-corrected reference genomes using dnadiff, the number of INDELs and SNV errors/100kb was calculated (Figure 4, Supplementary figure S 4, and Supplementary table S 1). Hybrid assemblies using R9.4.1+Illumina or R10.4.1+Illumina data had some of the lowest indel and SNV error rates, largely unaffected by long-read sequencing depth, but apparently differing by species (highest error rates for *E. coli* and lowest for *P. aeruginosa*; yellow and green lines, Figure 4). Long-read only assembly using R9.4.1 data (grey line, Figure 4) had the highest number of indel errors ranging between 1 and >10 indels/100kb, with some effect of increased sequencing depth in reducing this error rate; SNV error rates were generally lower than other long-read only assemblies, but only at sequencing depths >40x. R10.4.1 long-read only assemblies using data basecalled with the Dorado/rerio model (blue line, Figure 4) consistently had the lowest indel and SNV error rates amongst long-read only assemblies and for *E. coli* and *S. aureus* these were comparable to the hybrid assemblies.

**Figure 4.**
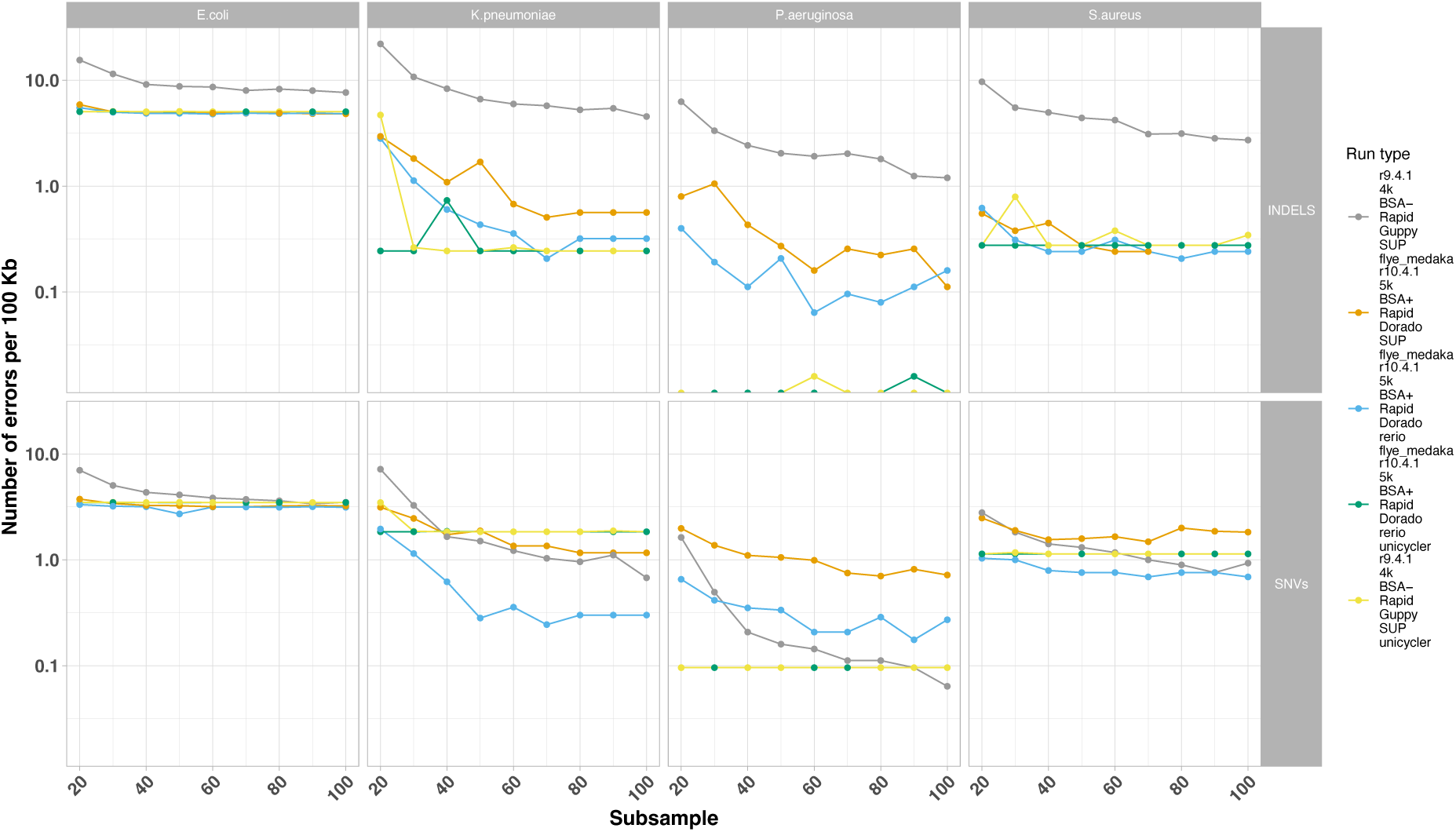
**Number of errors by error type (i.e. faceted row: single nucleotide [SNV]; insertion or deletion [INDEL]). X-axis represents subsampling to a specified depth (x coverage). Colours represent run type, including the flowcell cell used (R9.4.1 or R10.4.1), sampling rate (4k or 5k), if BSA was used (BSA+ or BSA-), the basecaller used (Guppy or dorado), the basecalling model used (SUP or rerio), and the assembly strategy used (Flye+Medaka or Unicycler) Unicycler assemblies represent nanopore-Illumina hybrid assemblies.**

Indel rates were notably highest for R9.4.1 long-read only assemblies, and predominantly biased to A and T deletions, except for *P. aeruginosa*, which notably has a high %GC content. These biases were less marked with R10.4.1 long-read only assemblies, with much lower indel rates. At a SNV level, G-to-A and C-to-T SNV errors appeared systematically more frequent across most sequencing/basecalling/assembly approaches and for most species; for R10.4.1 long-read only assemblies these were reduced by using the Dorado/rerio basecaller, except for the *E. coli* reference (Figure 5).

**Figure 5.**
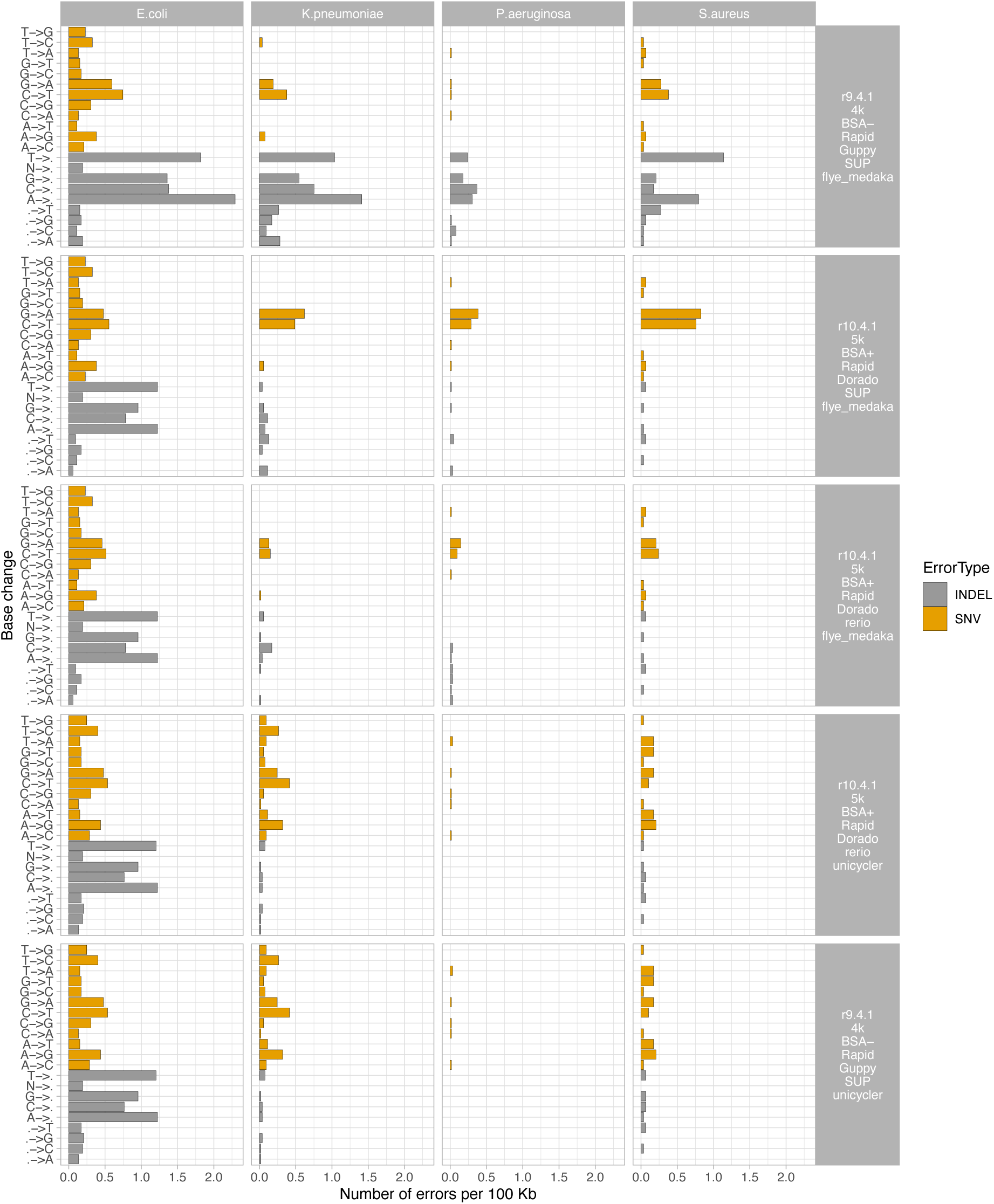
**Number of SNV and indel errors per 100 Kb by base change or gain/loss event (changes from reference to assembly), by species (faceted columns). Faceted rows represent run type, including the flowcell cell used (R9.4.1 or R10.4.1), sampling rate (4k or 5k), if BSA was used (BSA+ or BSA-), the basecaller used (Guppy or dorado), the basecalling model used (SUP or rerio), and the assembly strategy used (Flye+Medaka or Unicyler). Unicycler assemblies represent nanopore-Illumina hybrid assemblies.**

Compared to the coding sequences (CDS) annotated in the Illumina-corrected reference genomes, CDS content recovery was excellent for long-read only assemblies using R10.4.1 data and the Dorado/rerio model (i.e. above 99% at ≥30x coverage) and broadly comparable with gene content recovery for hybrid assemblies (Figure 6).

**Figure 6.**
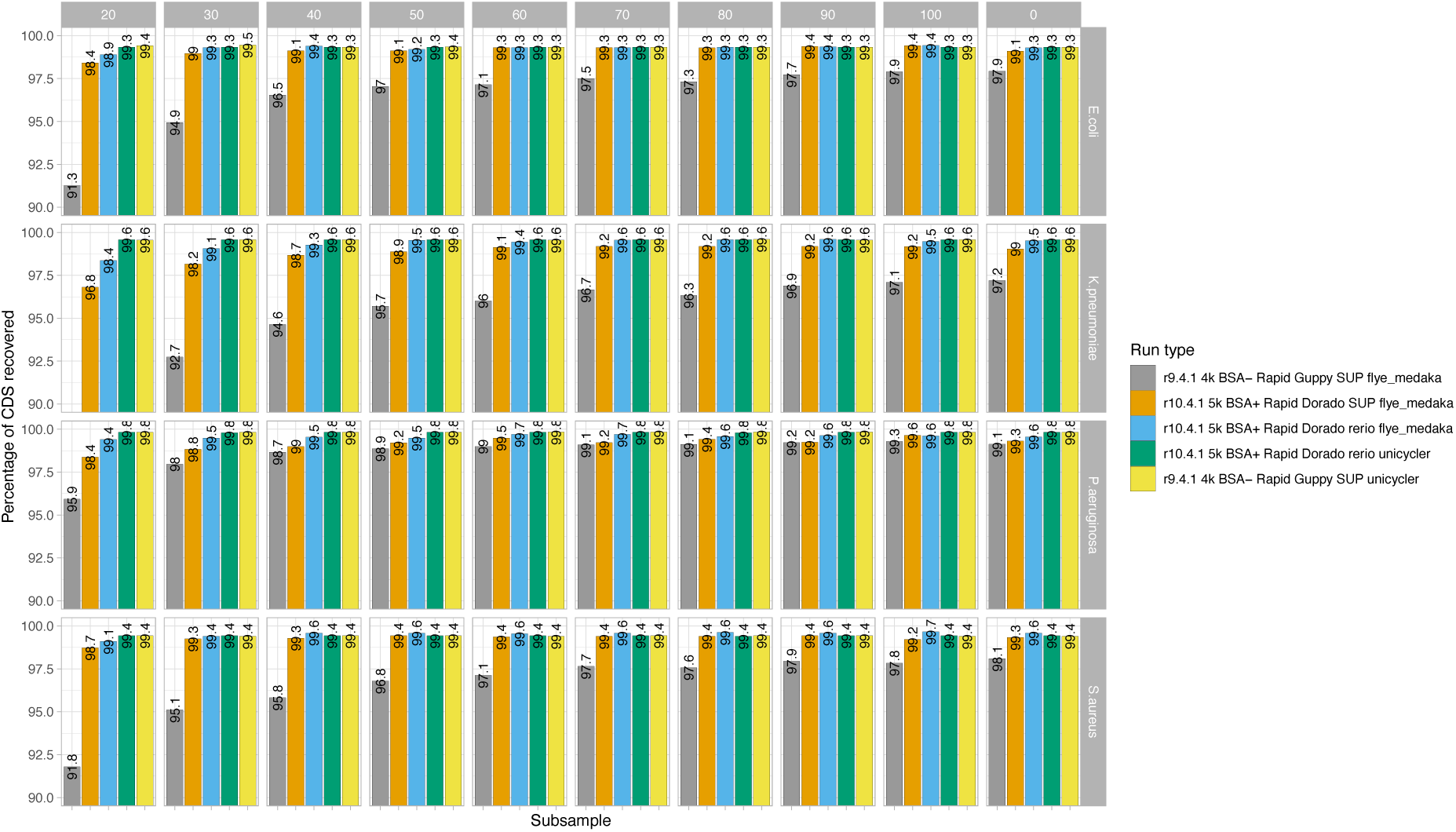
**Bar chart showing percentage of reference genome coding sequences (CDS) recovered by species (faceted rows) and subsampling to a specified depth (x coverage; faceted columns), with 0 the full number of reads from the run used. Colours represent run type, including the flowcell cell used (R9.4.1 or R10.4.1), sampling rate (4k or 5k), if BSA was used (BSA+ or BSA-), the basecaller used (Guppy or dorado), he basecalling model used (SUP or rerio)., and the assembly strategy used (Flye+Medaka or Unicycler). Unicycler assemblies represent nanopore-Illumina hybrid assemblies.**

#### 7.4.1 *E. coli* isolate sequencing - sequencing yield and read length distributions with and without BSA

The total sequencing yield for the run with BSA (BSA+) was 7.03 Gb, versus the run without BSA (BSA-) at 5.72 Gb. Following demultiplexing, ten isolates achieved enough bases for predicted ≥20x depth of coverage, however only nine empirically achieved this after mapping to the reference genomes and were used in further analysis (Supplementary figure S 5). Median (IQR) read lengths were shorter for the BSA+ compared to BSA-groups: 2.78 Kb (IQR: 0.81-6.75 Kb) versus 3.48 Kb respectively (IQR: 1.38-7.22 Kb; p<0.001; two-sample Wilcoxon); Supplementary figure S 6 and Supplementary figure S 7). However, BSA treatment resulted in greater per-read accuracy across sequenced *E. coli* isolates (BSA+ median %identity to the reference 99.04% compared to BSA-median % identity 98.22%; p <0.001, two sample Wilcoxon; Supplementary figure S 8).

R9.4.1+Illumina hybrid reference assemblies for the nine *E. coli* isolates included in the final analysis had chromosome sizes ranging from 4.7-5.26 Mb and contained between 2-8 plasmids, ranging in size from ∼1.5 Kb to ∼150 Kb (Table 2). Subsequently, using R10.4.1 long-read-only assembly, overall chromosome and plasmid recovery was roughly equal irrespective of the use of BSA, where incomplete assemblies were generated for some isolates and did not improve with increased depth (Figure 7). All chromosomes and plasmids were at least partially recovered at subsampling depths ≥20x. However, BSA+ outputs showed a reduction of indel and SNV error rates in 7/9 assembled genomes at ≥40x coverage (Figure 8).

**Figure 7.**
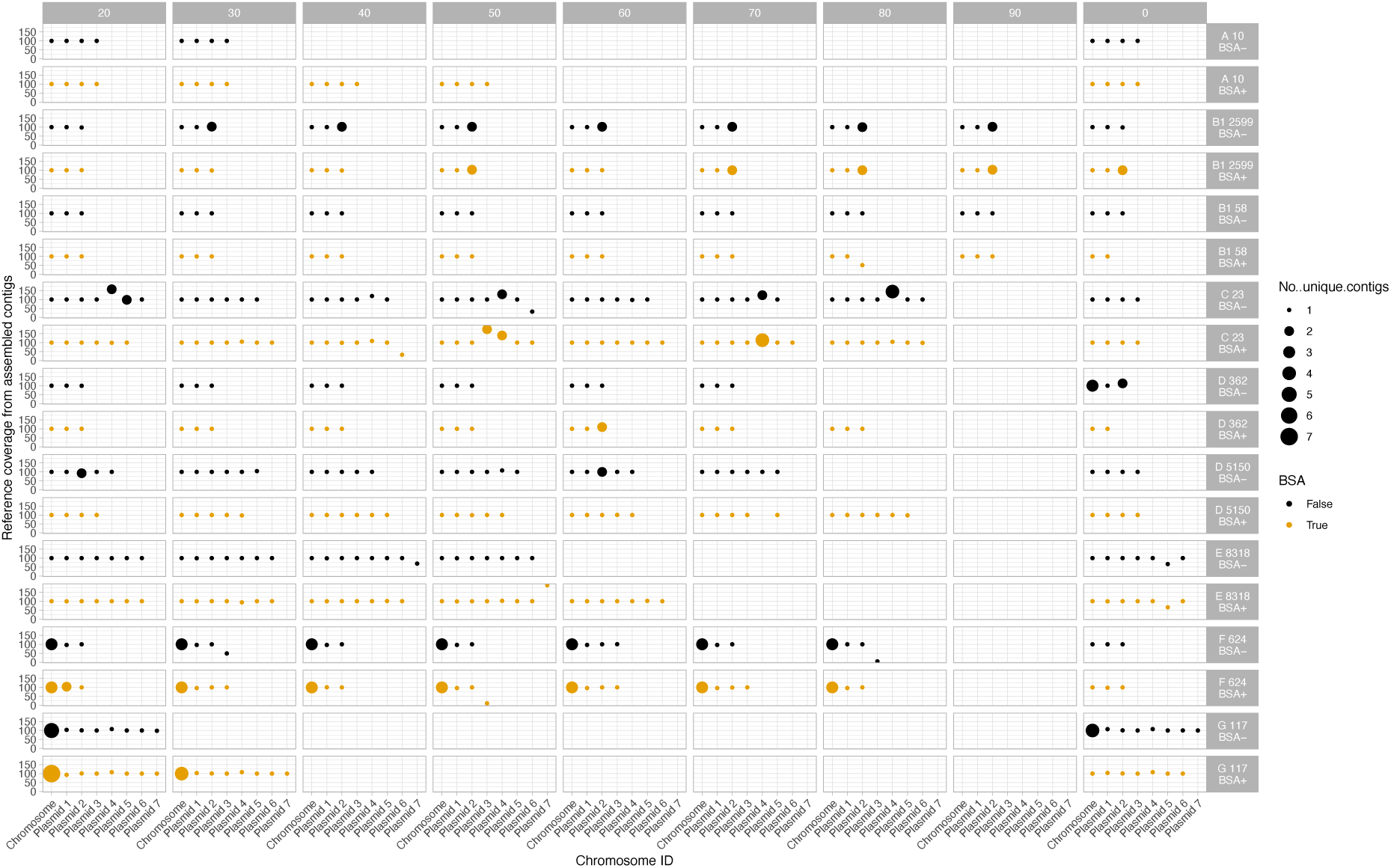
**Percentage of reference chromosome and plasmids recovered for each of nine sequenced clinical *E. coli* isolates passing QCwith (yellow) and without (black) BSA use in library treatment. Size of dots represent the number of unique contigs. Panels without any dots represent exclusions due to lack of coverage to subsample at this depth. The number of plasmids varies by isolate and plasmid number reflects the size order of plasmids within an isolate and does not correspond to the same plasmid between isolates. The letter number combination preceding the isolate numeric identifier indicates the *E. coli* phylogroup of the isolate.**

**Figure 8.**
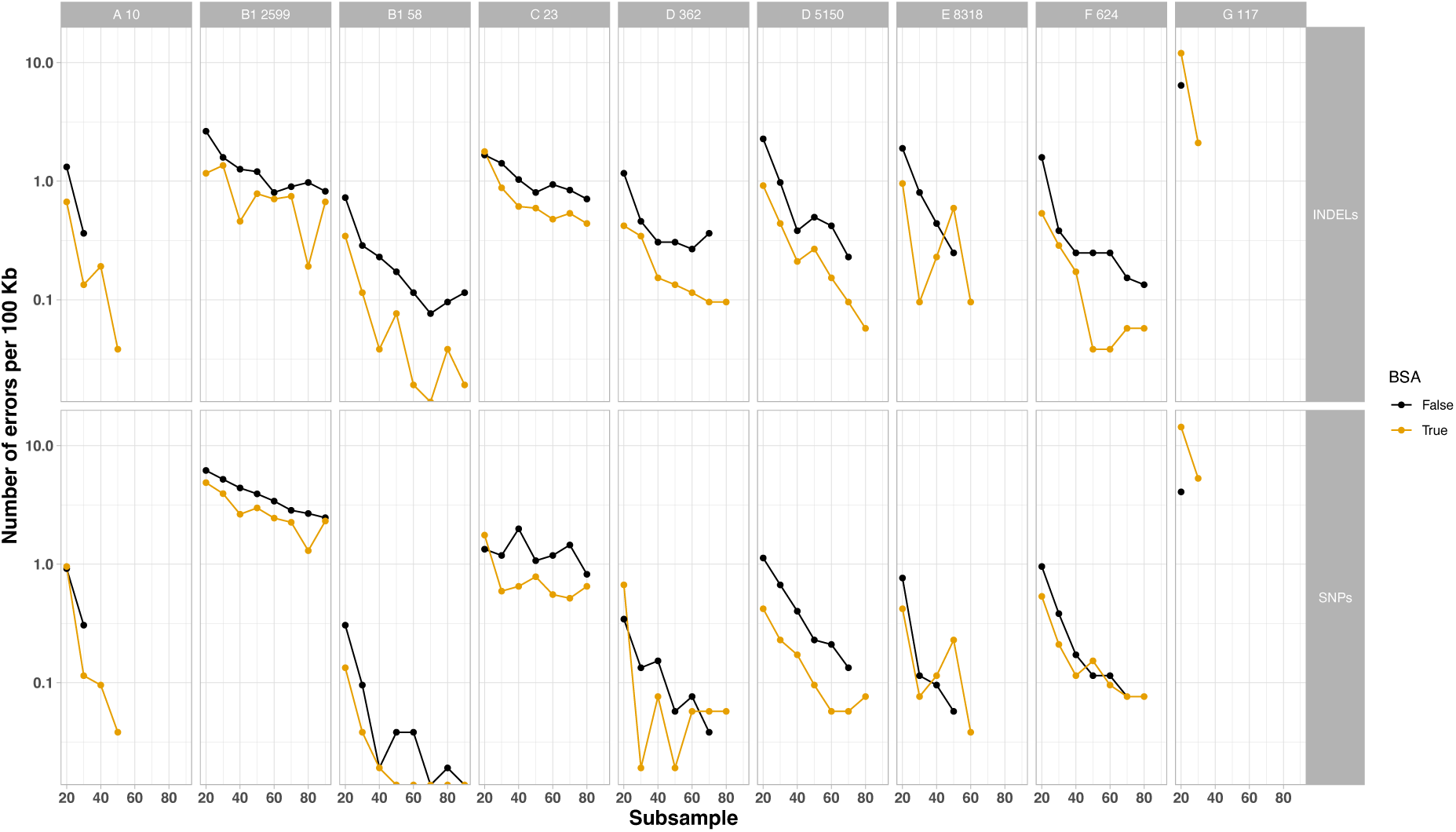
**Number of errors per 100kb over subsampled depth for each of nine sequenced clinical *E. coli* isolates passing QC with (yellow) and without (black) BSA use in library treatment. Rows show type, single nucleotide (SNV), insertion or deletion (INDEL). Faceted columns reflect different isolates. The letter number combination preceding the isolate numeric identifier indicates the *E. coli* phylogroup of the isolate.**

**Table 2.** ***E. coli* isolate reference chromosome and plasmid lengths (bp) from hybrid assemblies for each of nine sequenced clinical *E. coli* isolates. The letter number combination preceding the isolate numeric identifier indicates the *E. coli* phylogroup of the isolate.**

| Sample name | Chromosome | Plasmid 1 | Plasmid 2 | Plasmid 3 | Plasmid 4 | Plasmid 5 | Plasmid 6 | Plasmid 7 |
| --- | --- | --- | --- | --- | --- | --- | --- | --- |
| E 8318 | 4,838,664 | 93,041 | 71,835 | 10,877 | 5,772 | 3,371 | 2,096 | 1,552 |
| F 624 | 5,122,684 | 136,868 | 45,040 | 1,552 |  |  |  |  |
| D 362 | 5,094,307 | 137,038 | 4,671 |  |  |  |  |  |
| A 10 | 5,005,319 | 102,850 | 5,167 | 3,704 |  |  |  |  |
| C 23 | 4,701,962 | 145,252 | 99,455 | 46,065 | 2,328 | 2,101 | 1,545 |  |
| B2 131 | 5,007,009 | 83,691 | 69,217 | 6,082 | 5,167 | 4,087 | 1,822 | 1,565 |
| D 5150 | 5,258,227 | 138,055 | 48,768 | 5,166 | 2,678 | 2,090 |  |  |
| B1 58 | 5,127,313 | 151,663 | 4,715 |  |  |  |  |  |
| B1 2599 | 4,715,961 | 134,924 | 98,497 |  |  |  |  |  |
| G 117 | 5,164,489 | 140,103 | 115,805 | 59,734 | 10,579 | 7,119 | 5,114 | 4,072 |

## 8. Discussion

In this study we have shown that nanopore-only assemblies generated using R10.4.1 data to ∼40x depth, basecalling with Dorado and the updated Rerio model, and assembly with Flye and a single cycle of polishing with Medaka can be comparable to gold-standard Illumina-R9.4.1 or R10.4.1 nanopore hybrid assemblies (assembled using Unicycler). However, there appear to be species-specific differences (e.g. higher error rates for *E. coli*), and researchers should consider this carefully as part of research planning and use case. These may well reflect differences in methylation across strains and species, and are reduced by using the updated rerio model which has been trained with native bacterial DNA containing methylated sites[15]. Alternative strategies might include removing methylation from the bacterial DNA before sequencing using laboratory methods or a PCR-based library preparation - this would require further evaluation.

We also observed that using BSA during the library preparation generally improved sequencing yields and read accuracy, representing potential savings for sequencing costs and laboratory time. In our hands the R10.4.1 flowcells (BSA+) generate ∼6.4Gb using the rapid library kit. The consumables cost of such a sequencing run is currently approximately £1,200, resulting in a per isolate cost of £40-45 with 40x coverage (Table 3).

**Table 3.**
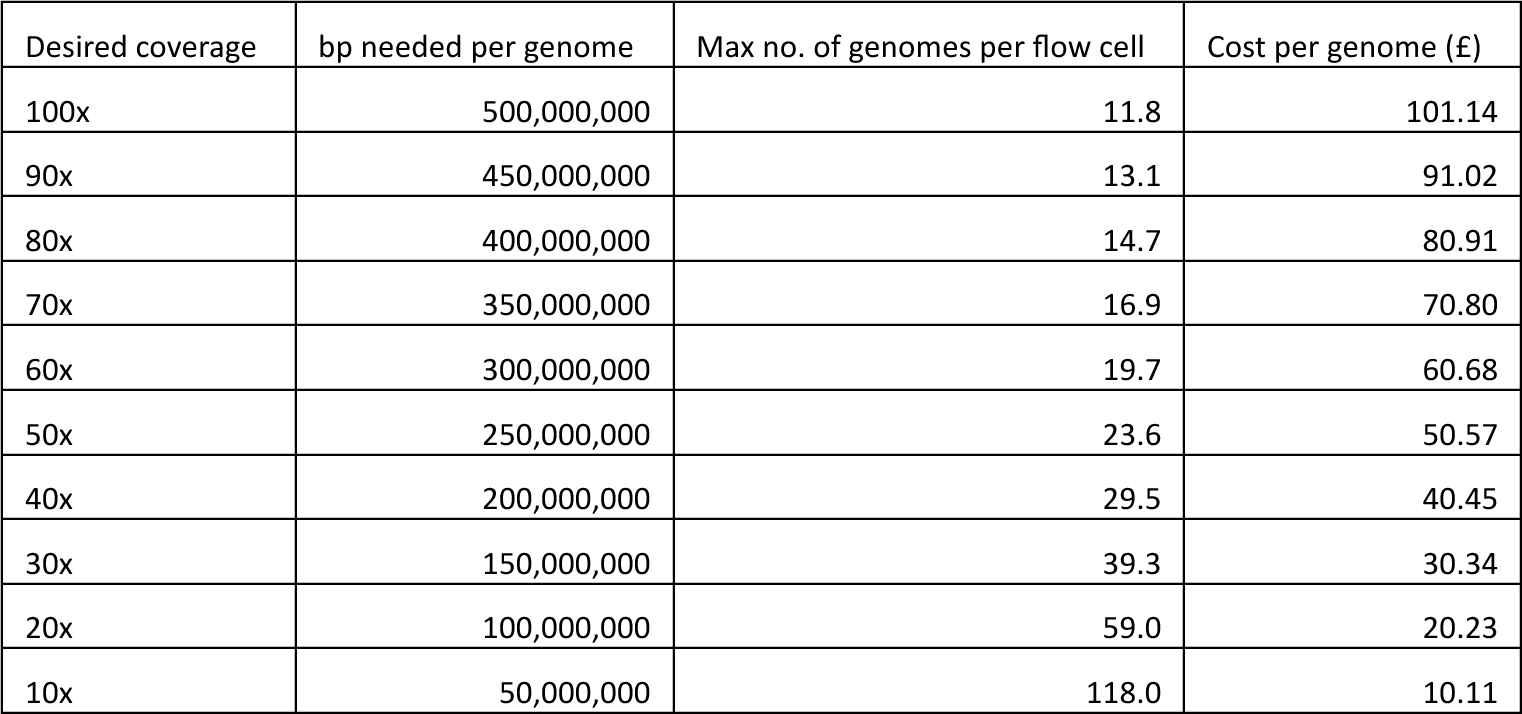
**Approximate R10.4.1-only sequencing cost per isolate at varying sequencing depths and genome sizes. Based on 5.9 Gb average yield from 3 R10.4.1 flowcells sequenced using the rapid library kit, and considering an approach using a 5k sampling rate, Dorado/Rerio basecalling and Flye assembly with one round of Medaka polishing.**

| Desired coverage | bp needed per genome | Max no. of genomes per flow cell | Cost per genome (£) |
| --- | --- | --- | --- |
| 100x | 500,000,000 | 11.8 | 101.14 |
| 90x | 450,000,000 | 13.1 | 91.02 |
| 80x | 400,000,000 | 14.7 | 80.91 |
| 70x | 350,000,000 | 16.9 | 70.80 |
| 60x | 300,000,000 | 19.7 | 60.68 |
| 50x | 250,000,000 | 23.6 | 50.57 |
| 40x | 200,000,000 | 29.5 | 40.45 |
| 30x | 150,000,000 | 39.3 | 30.34 |
| 20x | 100,000,000 | 59.0 | 20.23 |
| 10x | 50,000,000 | 118.0 | 10.11 |

### 8.1 Limitations

There are several limitations with this study. Our results are based on outputs from single runs only and there are no replicates for each sample and flowcell type; however, our findings of species-specific differences in sequencing outputs are similar to those observed by others[6, 16, 17]. Sequencing yield can be affected by numerous factors, including flowcell quality and baseline pore numbers, library preparation type, and operator skill. Replicating these findings by sequencing the same extracts using the same process across runs is important further work.

In this study we have compared annotated coding sequences between assembled genomes and the reference genome using a highly conservative requirement for a 100% match; gene presence/absence studies may require less stringent thresholds. Similarly, different use cases may tolerate higher indel/SNV error rates – as a result we have presented these without applying arbitrary acceptability thresholds.

Although we observed species-specific differences in error rates and assembly quality, we have only investigated single isolates from three species and 13 *E. coli* isolates. Our findings may not be generalisable across other strains and species, and we would advise that users consider characterising error rates and assembly quality for whichever species they are investigating by triangulating against a different method. We have not exhaustively investigated all possible combinations of sequencing, basecalling, assembly, and polishing strategy here, but have provided the raw data and reference assemblies to enable other users to evaluate their own workflows.

In the context of these limitations however, we have found that R10.4.1/Kit 14nanopore sequencing combined with updated, methylation-aware, basecalling models have improved bacterial genome reconstruction to enable reference quality genome assembly without the need for short-read sequencing and hybrid assembly. This could simplify pathogen genomics studies enabling cost-effective, higher throughput and faster turnaround times.

## 9. Figures and tables

**Supplementary figure S 1.**
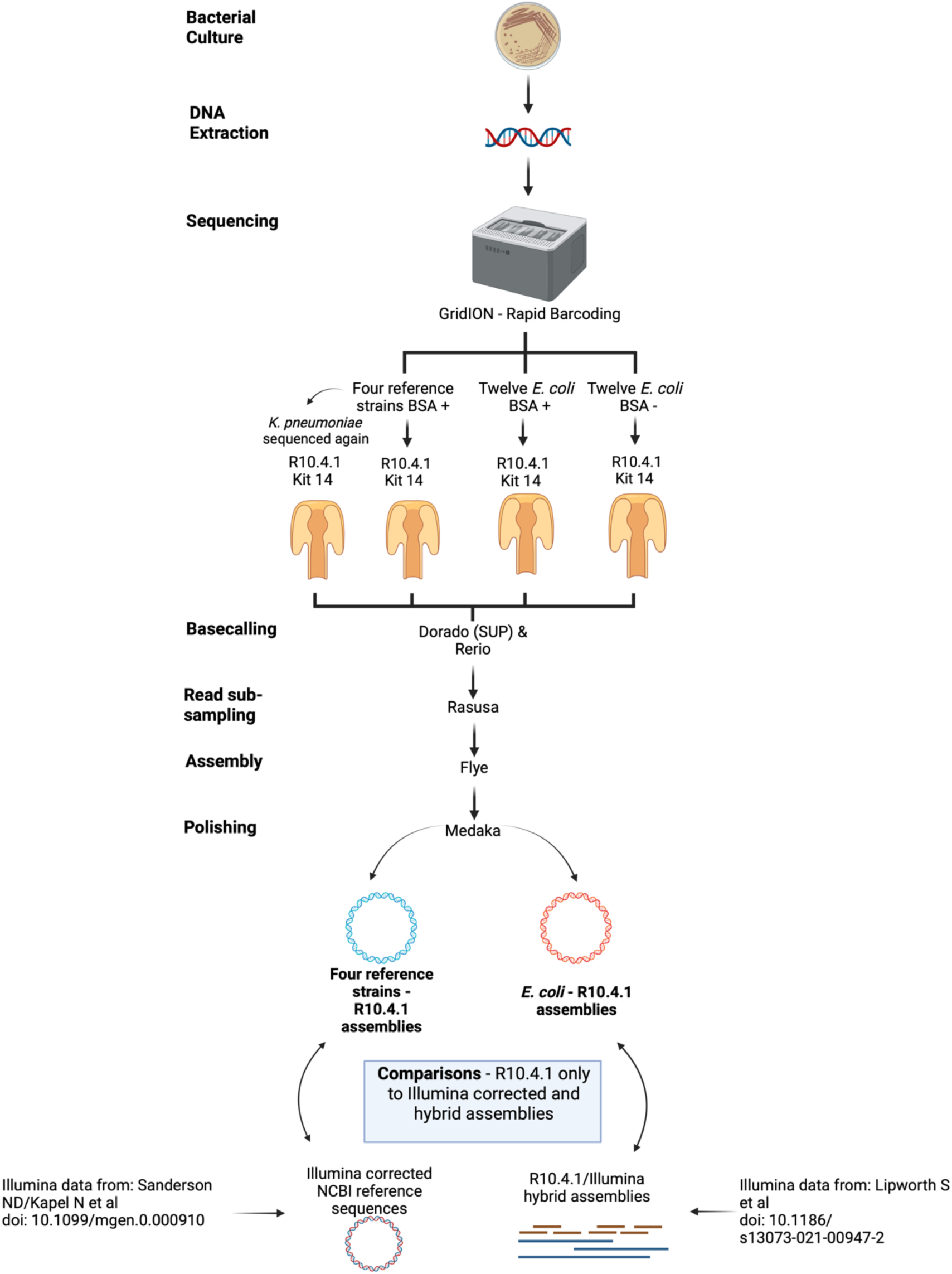
**Schematic of experimental workflow.**

**Supplementary figure S 2.**
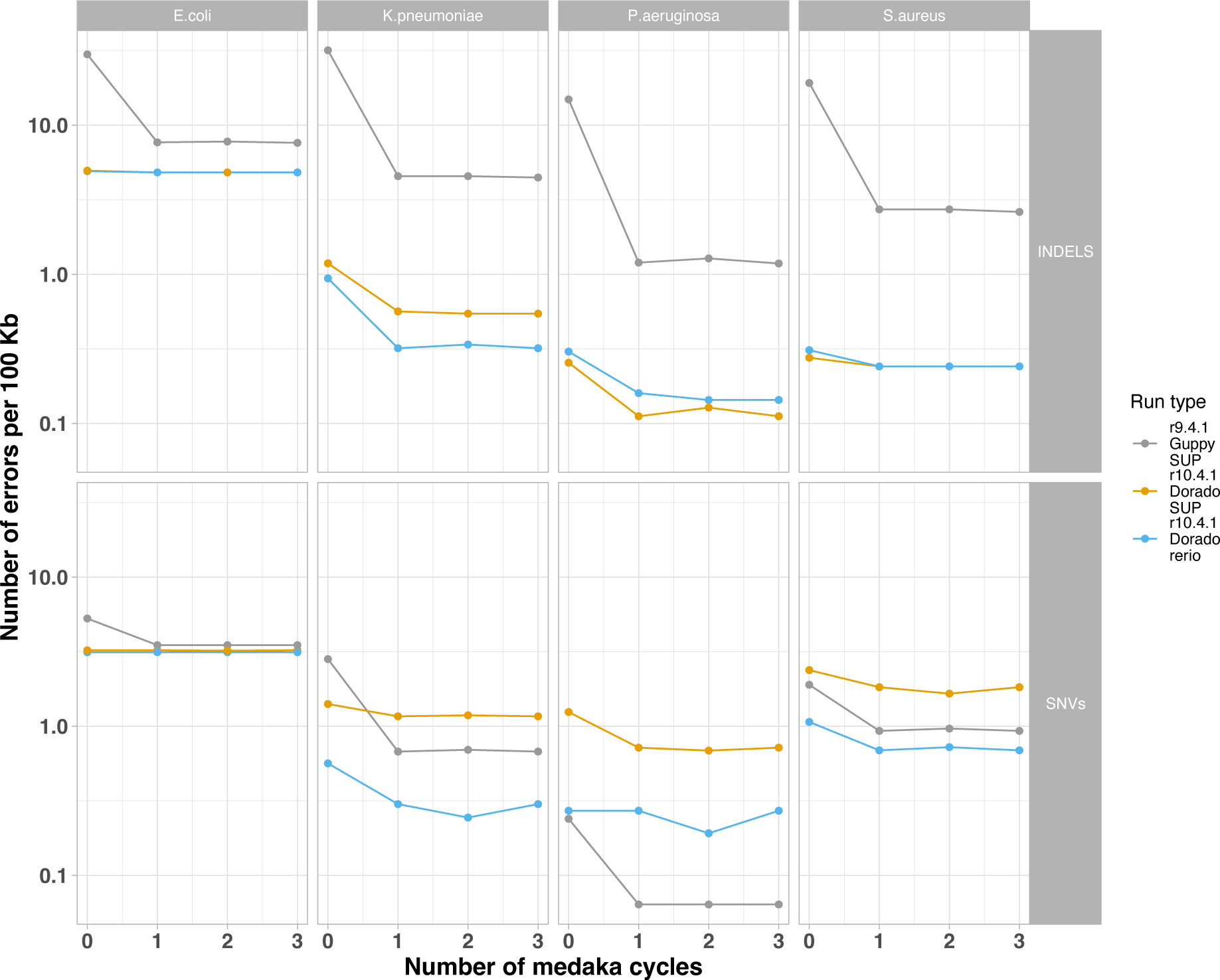
**Number of errors per 100kb by species and sequencing/basecalling approach using 1, 2, or 3 cycles of Medaka polishing.**

**Supplementary figure S 3.**
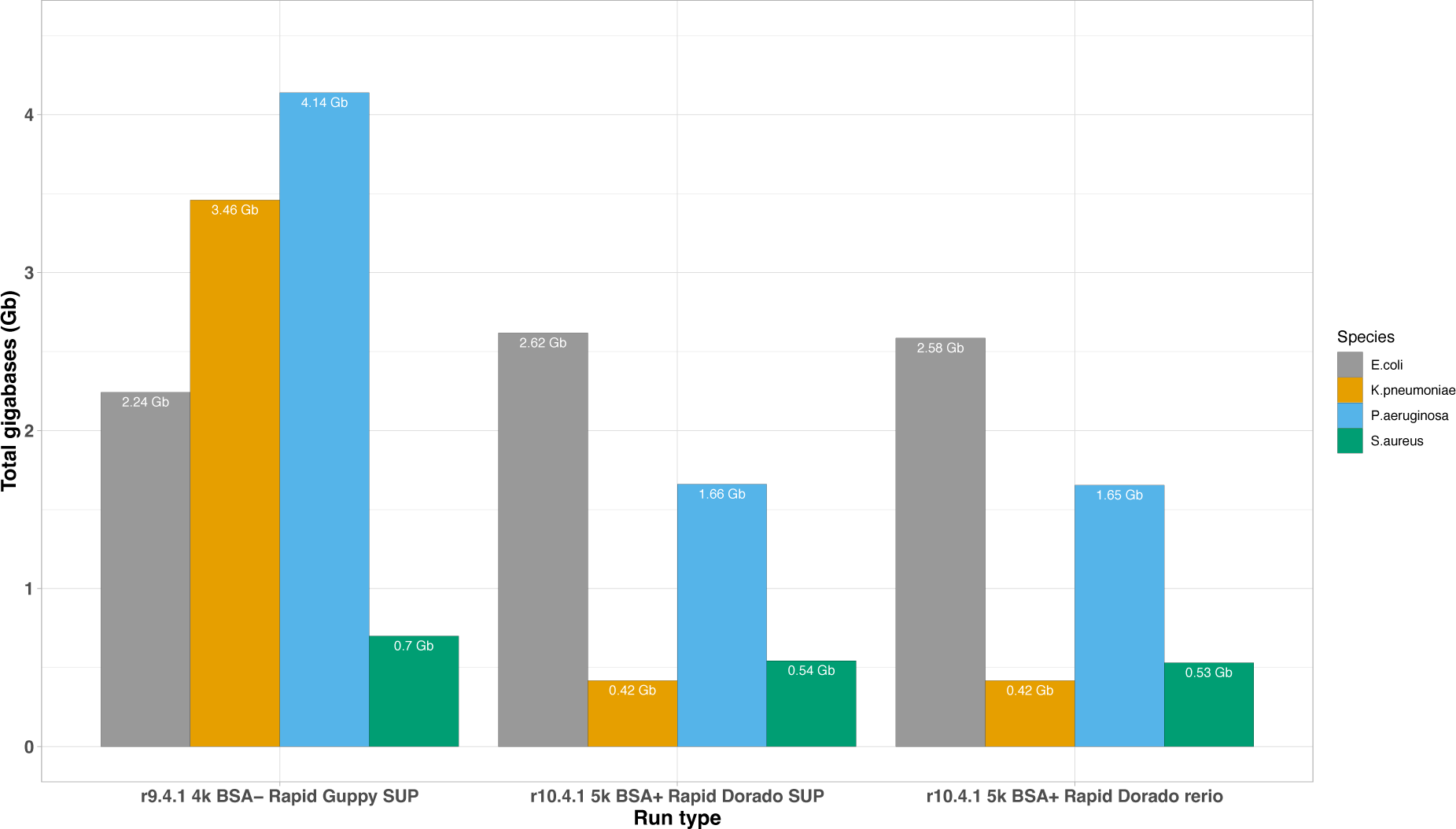
**Bar plot showing sequencing yield for reference strains sequenced, for each species by run type, including the flowcell cell used (R9.4.1 or R10.4.1), sampling rate (4k or 5k), if BSA was used (BSA+ or BSA-), the basecaller used (Guppy or dorado), and the basecalling model used (SUP or rerio).**

**Supplementary figure S 4.**
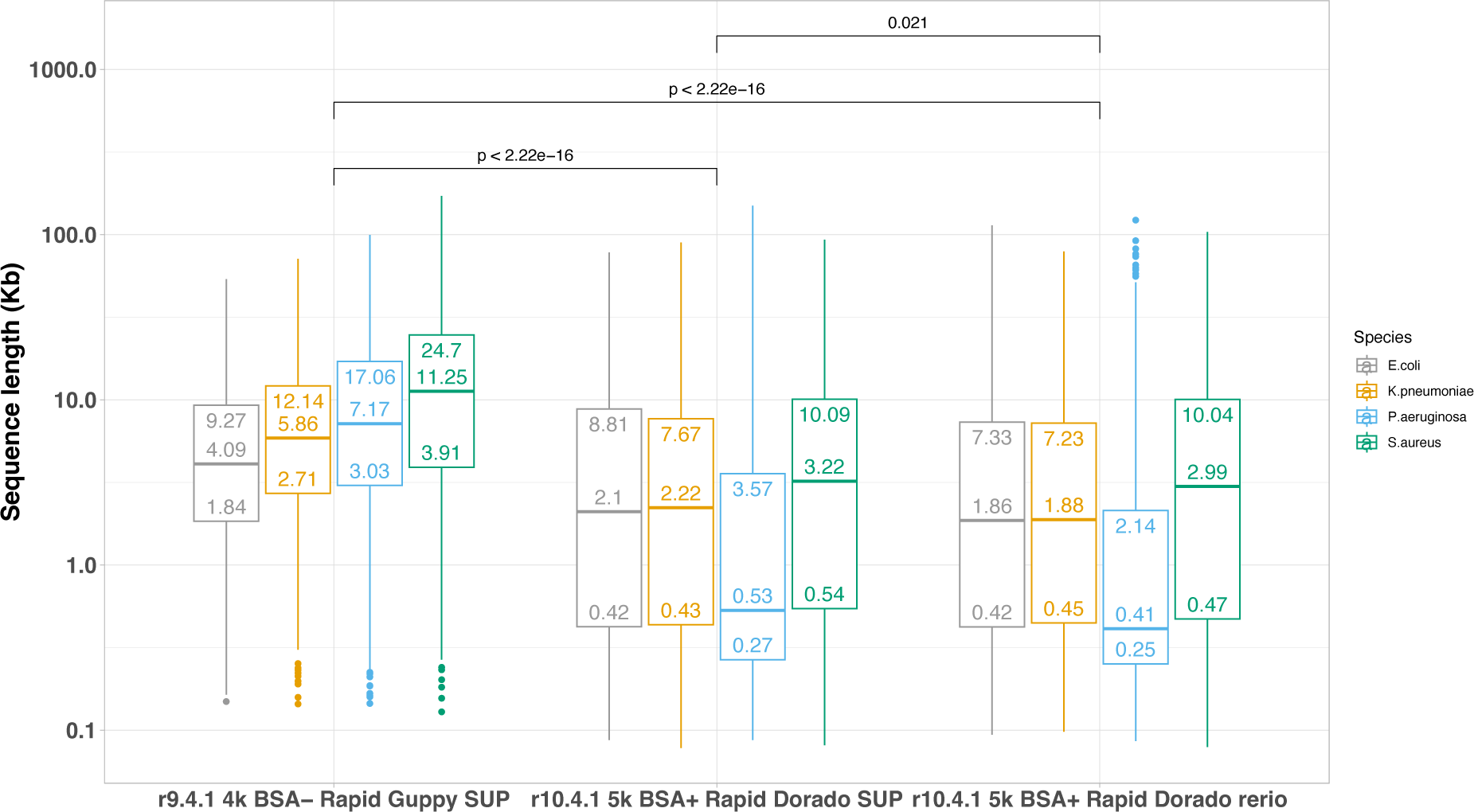
**Box and whisker plot of sequencing read lengths for reference strains sequenced, by sequencing run type and bacterial species. P-values calculated using two-sample Wilcoxon test.**

**Supplementary figure S 5.**
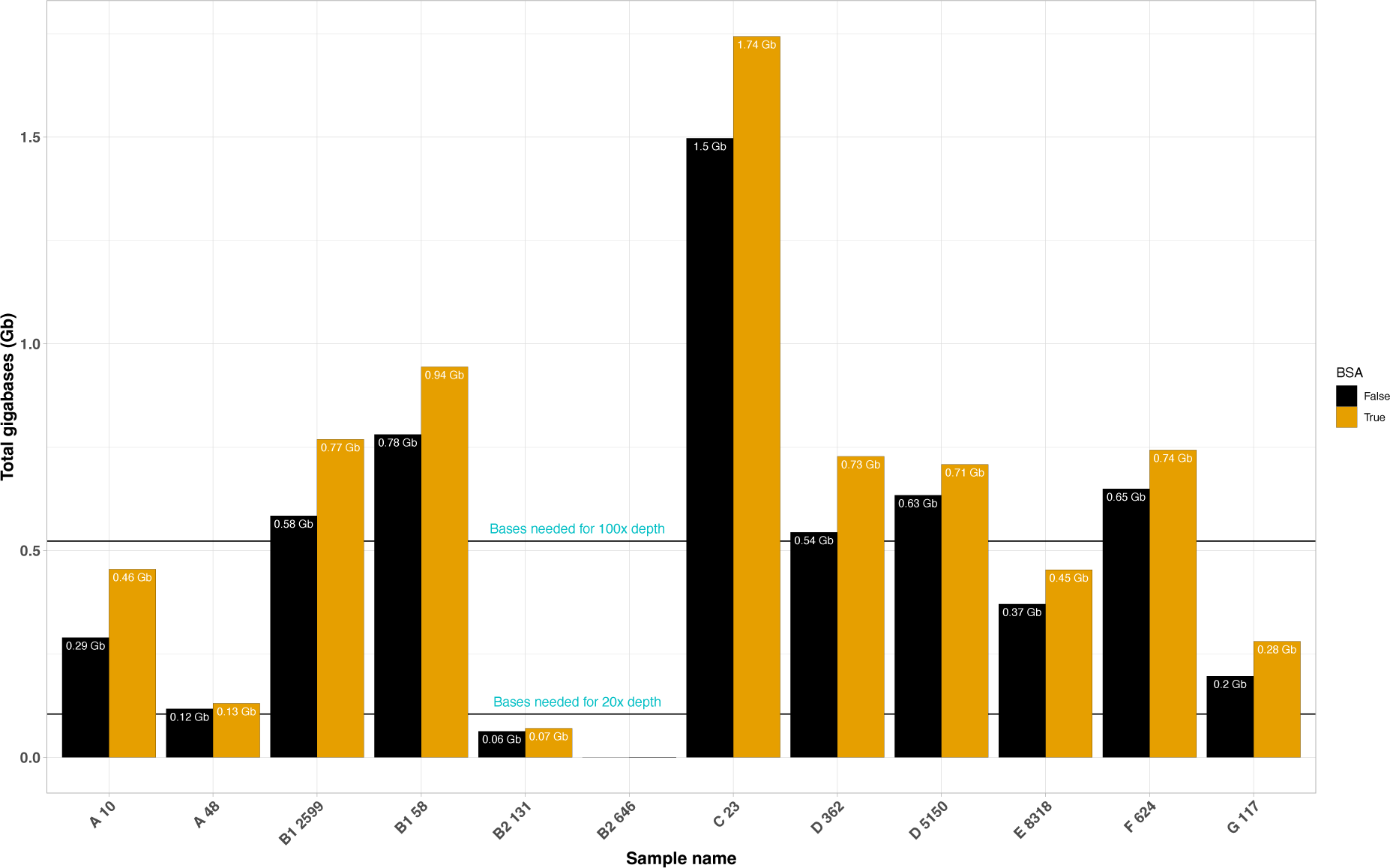
**Number of bases generated for each of twelve sequenced clinical *E. coli* isolates, with bovine serum albumin (BSA) (yellow) and without BSA used as part of library preparation (black). Lines showing number of bases required for theoretical depths of 20x and 100x coverage. The letter number combination preceding the isolate numeric identifier indicates the *E. coli* phylogroup of the isolate.**

**Supplementary figure S 6.**
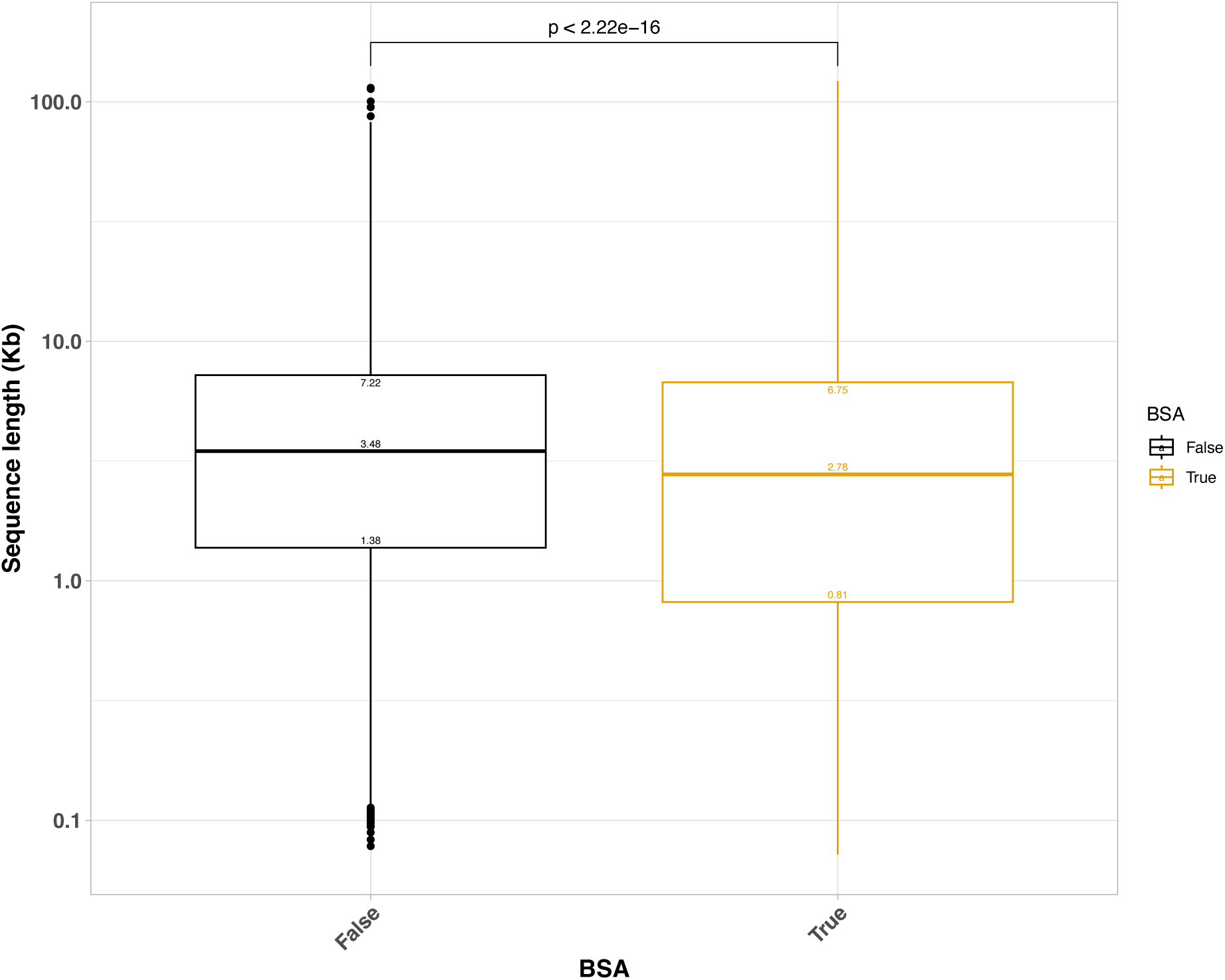
**Box and whisker plot of read lengths of 9 successful *E. coli* isolates sequenced with or without BSA. P-value calculated with two sample Wilcoxon test.**

**Supplementary figure S 7.**
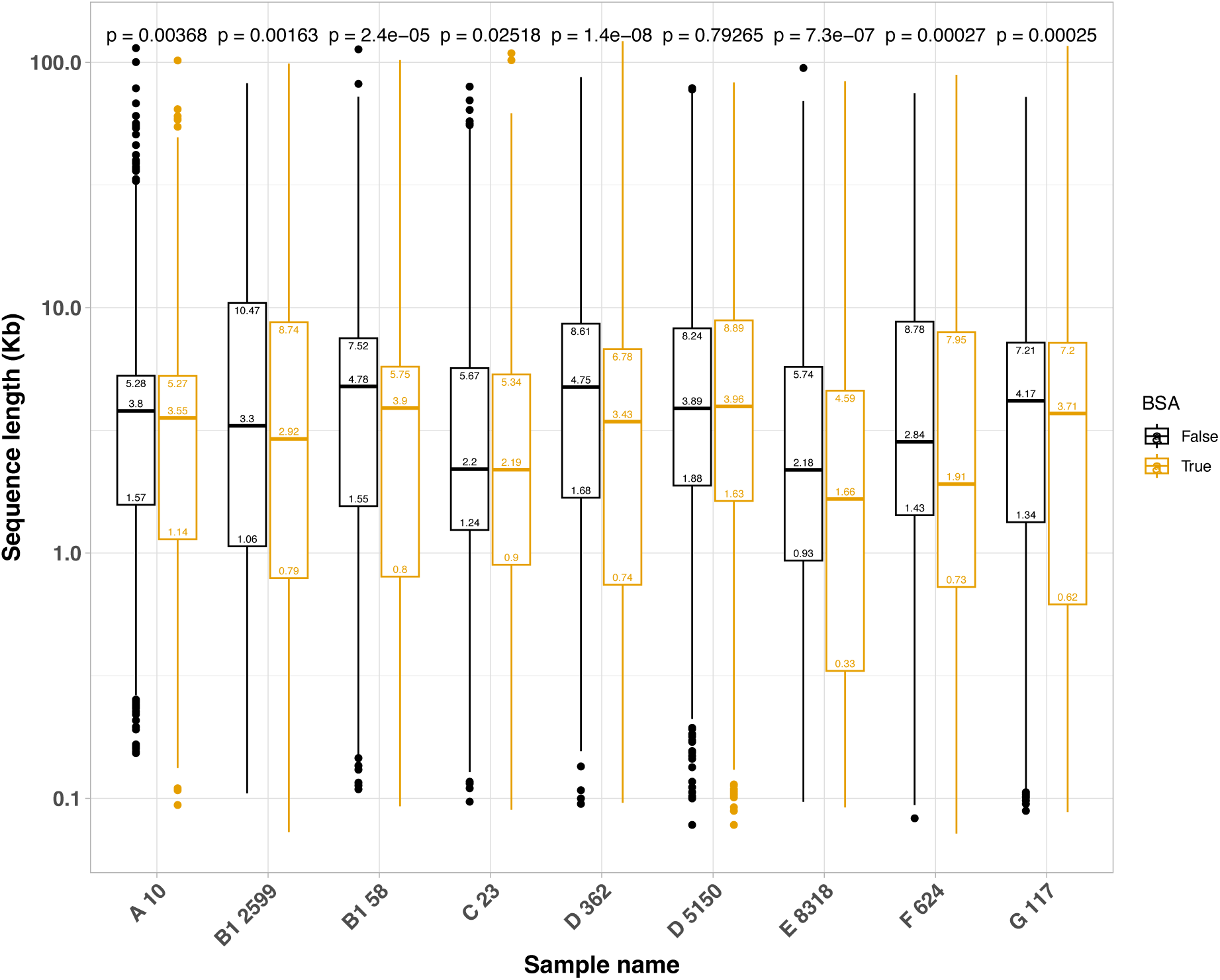
**Read length box and whisker plots for each of nine sequenced clinical *E. coli* isolates passing QC with (yellow) and without (black) BSA use in library preparation. P value calculated using an unpaired two sample Wilcoxon test. The letter number combination preceding the isolate numeric identifier indicates the *E. coli* phylogroup of the isolate.**

**Supplementary figure S 8.**
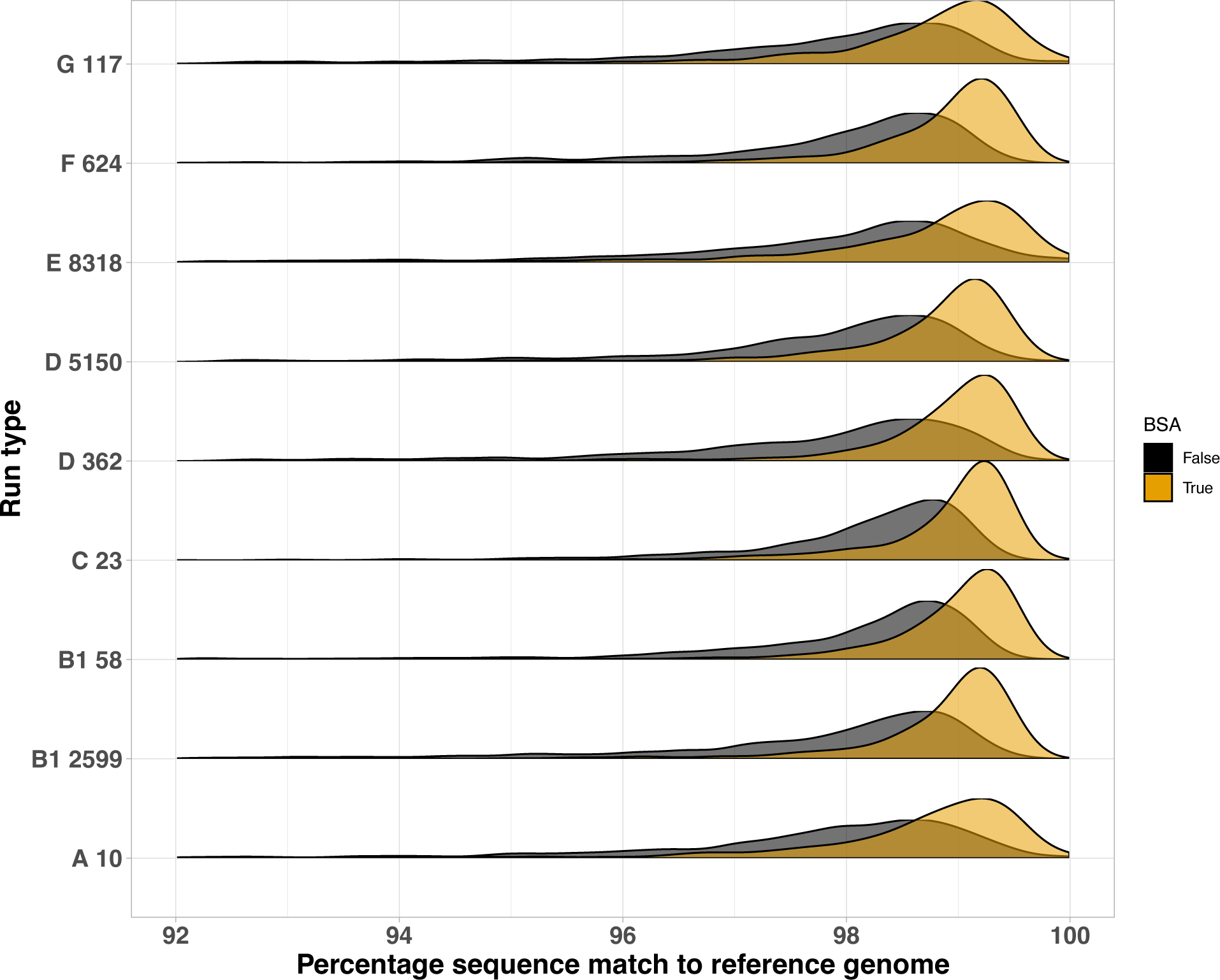
**Distributions of percentage read match to the reference genomes for each of nine sequenced clinical *E. coli* isolates passing QC with (yellow) and without (black) BSA use in library preparation. The letter number combination preceding the isolate numeric identifier indicates the *E. coli* phylogroup of the isolate.**

**Supplementary table S 1.** **Errors per 100 Kb for each flow cell run types at different subsampling depths.**

| Run.type | subsample | type | q1 | median | q3 |
| --- | --- | --- | --- | --- | --- |
| r10.4.1 Dorado rerio flye_medaka | 20 | INDELs | 0.5653575 | 1.72011926 | 3.49133469 |
| r10.4.1 Dorado rerio flye_medaka | 20 | SNVs | 0.93960009 | 1.49468499 | 2.29790538 |
| r10.4.1 Dorado rerio flye_medaka | 30 | INDELs | 0.28068194 | 0.71908219 | 2.0934746 |
| r10.4.1 Dorado rerio flye_medaka | 30 | SNVs | 0.85383387 | 1.07330827 | 1.66302168 |
| r10.4.1 Dorado rerio flye_medaka | 40 | INDELs | 0.20898975 | 0.42144153 | 1.67005707 |
| r10.4.1 Dorado rerio flye_medaka | 40 | SNVs | 0.55308499 | 0.7067021 | 1.38832663 |
| r10.4.1 Dorado rerio flye_medaka | 50 | INDELs | 0.23295142 | 0.33685507 | 1.54317737 |
| r10.4.1 Dorado rerio flye_medaka | 50 | SNVs | 0.32208617 | 0.54704197 | 1.24774181 |
| r10.4.1 Dorado rerio flye_medaka | 60 | INDELs | 0.24873306 | 0.33374384 | 1.46766594 |
| r10.4.1 Dorado rerio flye_medaka | 60 | SNVs | 0.31977408 | 0.55788177 | 1.35768445 |
| r10.4.1 Dorado rerio flye_medaka | 70 | INDELs | 0.17903685 | 0.22407311 | 1.40474385 |
| r10.4.1 Dorado rerio flye_medaka | 70 | SNVs | 0.23518761 | 0.46700804 | 1.30596031 |
| r10.4.1 Dorado rerio flye_medaka | 80 | INDELs | 0.17514046 | 0.26322271 | 1.44903068 |
| r10.4.1 Dorado rerio flye_medaka | 80 | SNVs | 0.29744889 | 0.52968628 | 1.35290433 |
| r10.4.1 Dorado rerio flye_medaka | 90 | INDELs | 0.20898975 | 0.28046409 | 1.46815114 |
| r10.4.1 Dorado rerio flye_medaka | 90 | SNVs | 0.26949362 | 0.52968628 | 1.36246456 |
| r10.4.1 Dorado rerio flye_medaka | 100 | INDELs | 0.22097058 | 0.28046409 | 1.44425056 |
| r10.4.1 Dorado rerio flye_medaka | 100 | SNVs | 0.29345528 | 0.49520353 | 1.30118019 |
| r10.4.1 Dorado rerio unicycler | 20 | INDELs | 0.18327068 | 0.26011149 | 1.46884684 |
| r10.4.1 Dorado rerio unicycler | 20 | SNVs | 0.87740994 | 1.49001815 | 2.25155983 |
| r10.4.1 Dorado rerio unicycler | 30 | INDELs | 0.18327068 | 0.26011149 | 1.46884684 |
| r10.4.1 Dorado rerio unicycler | 30 | SNVs | 0.87740994 | 1.49001815 | 2.25155983 |
| r10.4.1 Dorado rerio unicycler | 40 | INDELs | 0.20689655 | 0.50447239 | 1.81176232 |
| r10.4.1 Dorado rerio unicycler | 40 | SNVs | 0.87740994 | 1.49941665 | 2.26565757 |
| r10.4.1 Dorado rerio unicycler | 50 | INDELs | 0.18327068 | 0.26011149 | 1.46884684 |
| r10.4.1 Dorado rerio unicycler | 50 | SNVs | 0.87740994 | 1.49001815 | 2.25155983 |
| r10.4.1 Dorado rerio unicycler | 60 | INDELs | 0.18327068 | 0.26011149 | 1.46884684 |
| r10.4.1 Dorado rerio unicycler | 60 | SNVs | 0.87740994 | 1.49001815 | 2.25155983 |
| r10.4.1 Dorado rerio unicycler | 70 | INDELs | 0.18327068 | 0.26011149 | 1.46884684 |
| r10.4.1 Dorado rerio unicycler | 70 | SNVs | 0.87740994 | 1.49001815 | 2.25155983 |
| r10.4.1 Dorado rerio unicycler | 80 | INDELs | 0.18327068 | 0.26011149 | 1.46884684 |
| r10.4.1 Dorado rerio unicycler | 80 | SNVs | 0.87740994 | 1.49001815 | 2.25155983 |
| r10.4.1 Dorado rerio unicycler | 90 | INDELs | 0.18726429 | 0.26011149 | 1.46884684 |
| r10.4.1 Dorado rerio unicycler | 90 | SNVs | 0.87740994 | 1.49001815 | 2.25155983 |
| r10.4.1 Dorado rerio unicycler | 100 | INDELs | 0.18327068 | 0.26011149 | 1.46884684 |
| r10.4.1 Dorado rerio unicycler | 100 | SNVs | 0.87740994 | 1.49001815 | 2.25155983 |
| r10.4.1 Dorado SUP flye_medaka | 20 | INDELs | 0.73697257 | 1.87492493 | 3.6856212 |
| r10.4.1 Dorado SUP flye_medaka | 20 | SNVs | 2.35727663 | 2.81092818 | 3.29122579 |
| r10.4.1 Dorado SUP flye_medaka | 30 | INDELs | 0.88556241 | 1.43881068 | 2.62465138 |
| r10.4.1 Dorado SUP flye_medaka | 30 | SNVs | 1.76586427 | 2.17947887 | 2.69766493 |
| r10.4.1 Dorado SUP flye_medaka | 40 | INDELs | 0.44403437 | 0.76925071 | 2.07005923 |
| r10.4.1 Dorado SUP flye_medaka | 40 | SNVs | 1.43935221 | 1.64052372 | 2.1143921 |
| r10.4.1 Dorado SUP flye_medaka | 50 | INDELs | 0.27478793 | 0.9837957 | 2.51640694 |
| r10.4.1 Dorado SUP flye_medaka | 50 | SNVs | 1.45323345 | 1.73295307 | 2.22239394 |
| r10.4.1 Dorado SUP flye_medaka | 60 | INDELs | 0.22097058 | 0.45903552 | 1.73122817 |
| r10.4.1 Dorado SUP flye_medaka | 60 | SNVs | 1.26264143 | 1.50427794 | 2.03487835 |
| r10.4.1 Dorado SUP flye_medaka | 70 | INDELs | 0.25203812 | 0.38155493 | 1.59956835 |
| r10.4.1 Dorado SUP flye_medaka | 70 | SNVs | 1.20273727 | 1.41807104 | 1.90556801 |
| r10.4.1 Dorado SUP flye_medaka | 80 | INDELs | 0.21945577 | 0.39377597 | 1.6466417 |
| r10.4.1 Dorado SUP flye_medaka | 80 | SNVs | 1.049779 | 1.58270677 | 2.30783939 |
| r10.4.1 Dorado SUP flye_medaka | 90 | INDELs | 0.25203812 | 0.40975041 | 1.63230136 |
| r10.4.1 Dorado SUP flye_medaka | 90 | SNVs | 1.07773427 | 1.51374125 | 2.20917123 |
| r10.4.1 Dorado SUP flye_medaka | 100 | INDELs | 0.20898975 | 0.40264454 | 1.62752124 |
| r10.4.1 Dorado SUP flye_medaka | 100 | SNVs | 1.05377261 | 1.49649987 | 2.17852904 |
| r9.4.1 Guppy SUP flye_medaka | 20 | INDELs | 8.8367302 | 12.6077339 | 17.1612768 |
| r9.4.1 Guppy SUP flye_medaka | 20 | SNVs | 2.50217583 | 4.90515593 | 7.05801909 |
| r9.4.1 Guppy SUP flye_medaka | 30 | INDELs | 4.97259557 | 8.14395904 | 10.9508565 |
| r9.4.1 Guppy SUP flye_medaka | 30 | SNVs | 1.49449157 | 2.54913145 | 3.71495781 |
| r9.4.1 Guppy SUP flye_medaka | 40 | INDELs | 4.33116669 | 6.64629246 | 8.53019559 |
| r9.4.1 Guppy SUP flye_medaka | 40 | SNVs | 1.11226176 | 1.53396422 | 2.32568755 |
| r9.4.1 Guppy SUP flye_medaka | 50 | INDELs | 3.82152694 | 5.52456572 | 7.1657963 |
| r9.4.1 Guppy SUP flye_medaka | 50 | SNVs | 1.02269472 | 1.40705211 | 2.15554421 |
| r9.4.1 Guppy SUP flye_medaka | 60 | INDELs | 3.63440564 | 5.09217008 | 6.64369456 |
| r9.4.1 Guppy SUP flye_medaka | 60 | SNVs | 0.91525284 | 1.19710915 | 1.87715644 |
| r9.4.1 Guppy SUP flye_medaka | 70 | INDELs | 2.83477471 | 4.42766399 | 6.31677784 |
| r9.4.1 Guppy SUP flye_medaka | 70 | SNVs | 0.77795527 | 1.01691729 | 1.70749831 |
| r9.4.1 Guppy SUP flye_medaka | 80 | INDELs | 2.80472623 | 4.20054446 | 6.01237798 |
| r9.4.1 Guppy SUP flye_medaka | 80 | SNVs | 0.70036906 | 0.92759917 | 1.62242665 |
| r9.4.1 Guppy SUP flye_medaka | 90 | INDELs | 2.43219125 | 4.12995852 | 6.07233607 |
| r9.4.1 Guppy SUP flye_medaka | 90 | SNVs | 0.59292718 | 0.93382162 | 1.67784722 |
| r9.4.1 Guppy SUP flye_medaka | 100 | INDELs | 2.34262422 | 3.63650506 | 5.32848014 |
| r9.4.1 Guppy SUP flye_medaka | 100 | SNVs | 0.52349324 | 0.80386311 | 1.57303686 |
| r9.4.1 Guppy SUP unicycler | 20 | INDELs | 0.20689655 | 2.48755509 | 4.78638638 |
| r9.4.1 Guppy SUP unicycler | 20 | SNVs | 0.87740994 | 2.30768732 | 3.4828437 |
| r9.4.1 Guppy SUP unicycler | 30 | INDELs | 0.19736842 | 0.52813067 | 1.8663381 |
| r9.4.1 Guppy SUP unicycler | 30 | SNVs | 0.90327201 | 1.51665802 | 2.27043769 |
| r9.4.1 Guppy SUP unicycler | 40 | INDELs | 0.18327068 | 0.26011149 | 1.46884684 |
| r9.4.1 Guppy SUP unicycler | 40 | SNVs | 0.87740994 | 1.49001815 | 2.25155983 |
| r9.4.1 Guppy SUP unicycler | 50 | INDELs | 0.18327068 | 0.26011149 | 1.47840707 |
| r9.4.1 Guppy SUP unicycler | 50 | SNVs | 0.87740994 | 1.49001815 | 2.25633994 |
| r9.4.1 Guppy SUP unicycler | 60 | INDELs | 0.20136203 | 0.32123412 | 1.54643305 |
| r9.4.1 Guppy SUP unicycler | 60 | SNVs | 0.87740994 | 1.49001815 | 2.25155983 |
| r9.4.1 Guppy SUP unicycler | 70 | INDELs | 0.18327068 | 0.26011149 | 1.46884684 |
| r9.4.1 Guppy SUP unicycler | 70 | SNVs | 0.87740994 | 1.49001815 | 2.25155983 |
| r9.4.1 Guppy SUP unicycler | 80 | INDELs | 0.18327068 | 0.26011149 | 1.46884684 |
| r9.4.1 Guppy SUP unicycler | 80 | SNVs | 0.87740994 | 1.49001815 | 2.25155983 |
| r9.4.1 Guppy SUP unicycler | 90 | INDELs | 0.18327068 | 0.26011149 | 1.46884684 |
| r9.4.1 Guppy SUP unicycler | 90 | SNVs | 0.87740994 | 1.50881514 | 2.27975532 |
| r9.4.1 Guppy SUP unicycler | 100 | INDELs | 0.18327068 | 0.29459424 | 1.52057098 |
| r9.4.1 Guppy SUP unicycler | 100 | SNVs | 0.87740994 | 1.49001815 | 2.25155983 |

## 10. Author statements

### 10.1 Author contributions

KH, NDS and NS designed the study. KH and NK performed the laboratory experiments and sequencing. NDS performed the bioinformatics analysis. NSt, NDS and KH wrote the first draft. All authors reviewed and approved the final draft.

### 10.4 Ethical approval

Not applicable.

### 10.5 Consent for publication

Not applicable.

## 10.6 Acknowledgements

We are grateful to Dr Celiq Souque and Prof Craig Maclean at the Department of Zoology, University of Oxford, for supplying the *Pseudomonas aeruginosa* PAO1 strain.

## References

1. Baker KS, Jauneikaite E, Hopkins KL, Lo SW, Sánchez-Busó L, et al. Genomics for public health and international surveillance of antimicrobial resistance. The Lancet Microbe 2023;4:e1047–e1055.

2. De Maio N, Shaw LP, Hubbard A, George S, Sanderson ND, et al. Comparison of long-read sequencing technologies in the hybrid assembly of complex bacterial genomes. Microb Genom;5.

3. Accuracy. *Oxford Nanopore Technologies*. https://nanoporetech.com/accuracy (accessed 12 January 2024).

4. Sanderson ND, Kapel N, Rodger G, Webster H, Lipworth S, et al. Comparison of R9.4.1/Kit10 and R10/Kit12 Oxford Nanopore flowcells and chemistries in bacterial genome reconstruction. Microbial Genomics 2023;9:000910.

5. Kirkegaard R. Kirk3gaard/2023-basecalling-benchmarks. https://github.com/Kirk3gaard/2023-basecalling-benchmarks (2023, accessed 19 December 2023).

6. Wick R. ONT-only accuracy with R10.4.1. Ryan Wick’s bioinformatics blog. https://rrwick.github.io/2023/05/05/ont-only-accuracy-with-r10.4.1.html (2023, accessed 19 December 2023).

7. Lipworth S, Vihta K-D, Chau K, Barker L, George S, et al. Ten-year longitudinal molecular epidemiology study of Escherichia coli and Klebsiella species bloodstream infections in Oxfordshire, UK. Genome Med 2021;13:144.

8. Beghain J, Bridier-Nahmias A, Le Nagard H, Denamur E, Clermont O. ClermonTyping: an easy-to-use and accurate in silico method for Escherichia genus strain phylotyping. Microb Genom 2018;4:e000192.

9. Wick RR, Judd LM, Gorrie CL, Holt KE. Unicycler: Resolving bacterial genome assemblies from short and long sequencing reads. PLOS Computational Biology 2017;13:e1005595.

10. Hall MB. Rasusa: Randomly subsample sequencing reads to a specified coverage. Journal of Open Source Software 2022;7:3941.

11. Kolmogorov M, Yuan J, Lin Y, Pevzner PA. Assembly of long, error-prone reads using repeat graphs. Nat Biotechnol 2019;37:540–546.

12. Li H. Minimap2: fast pairwise alignment for long nucleotide sequences. 2017;2–5.

13. Li H, Handsaker B, Wysoker A, Fennell T, Ruan J, et al. The Sequence Alignment/Map format and SAMtools. Bioinformatics (Oxford, England) 2009;25:2078–2079.

14. Kurtz S, Phillippy A, Delcher AL, Smoot M, Shumway M, et al. Versatile and open software for comparing large genomes. Genome Biology.

15. Community. Oxford Nanopore Technologies. https://community.nanoporetech.com/posts/ligation-sequencing-kit-v1 (accessed 19 December 2023).

16. Wick R. ONT-only accuracy: 5 kHz and Dorado. Ryan Wick’s bioinformatics blog. https://rrwick.github.io/2023/10/24/ont-only-accuracy-update.html (2023, accessed 20 December 2023).

17. Sereika M, Kirkegaard RH, Karst SM, Michaelsen TY, Sørensen EA, et al. Oxford Nanopore R10.4 long-read sequencing enables the generation of near-finished bacterial genomes from pure cultures and metagenomes without short-read or reference polishing. Nat Methods 2022;19:823–826.

